# Binding specificities of human RNA binding proteins towards structured and linear RNA sequences

**DOI:** 10.1101/317909

**Authors:** Arttu Jolma, Jilin Zhang, Estefania Mondragón, Ekaterina Morgunova, Teemu Kivioja, Kaitlin U. Laverty, Yimeng Yin, Fangjie Zhu, Gleb Bourenkov, Quaid Morris, Timothy R. Hughes, Louis James Maher, Jussi Taipale

**Affiliations:** Department of Medical Biochemistry and Biophysics, Karolinska Institutet, Solna, Sweden; Department of Biochemistry and Molecular Biology and Mayo Clinic Graduate School of Biomedical Sciences, Mayo Clinic College of Medicine and Science, Rochester, USA; Genome-Scale Biology Program, University of Helsinki, Helsinki, Finland; Department of Molecular Genetics, University of Toronto, Toronto, Canada; European Molecular Biology Laboratory (EMBL), Hamburg Unit c/o DESY, Notkestrasse 85, D-22603 Hamburg, Germany; Donnelly Centre, University of Toronto, Toronto, Canada; Edward S Rogers Sr Department of Electrical and Computer Engineering, University of Toronto, Toronto, Canada; Department of Computer Science, University of Toronto, Toronto, Canada; Department of Biochemistry, University of Cambridge, Cambridge, United Kingdom

## Abstract

Sequence specific RNA-binding proteins (RBPs) control many important processes affecting gene expression. They regulate RNA metabolism at multiple levels, by affecting splicing of nascent transcripts, RNA folding, base modification, transport, localization, translation and stability. Despite their central role in most aspects of RNA metabolism and function, most RBP binding specificities remain unknown or incompletely defined. To address this, we have assembled a genome-scale collection of RBPs and their RNA binding domains (RBDs), and assessed their specificities using high throughput RNA-SELEX (HTR-SELEX). Approximately 70% of RBPs for which we obtained a motif bound to short linear sequences, whereas ~30% preferred structured motifs folding into stem-loops. We also found that many RBPs can bind to multiple distinctly different motifs. Analysis of the matches of the motifs in human genomic sequences suggested novel roles for many RBPs. We found that three cytoplasmic proteins, ZC3H12A, ZC3H12B and ZC3H12C bound to motifs resembling the splice donor sequence, suggesting that these proteins are involved in degradation of cytoplasmic viral and/or unspliced transcripts. Surprisingly, structural analysis revealed that the RNA motif was not bound by the conventional C3H1 RNA-binding domain of ZC3H12B. Instead, the RNA motif was bound by the ZC3H12B’s PilT N-terminus (PIN) RNase domain, revealing a potential mechanism by which unconventional RNA binding domains containing active sites or molecule-binding pockets could interact with short, structured RNA molecules. Our collection containing 145 high resolution binding specificity models for 86 RBPs is the largest systematic resource for the analysis of human RBPs, and will greatly facilitate future analysis of the various biological roles of this important class of proteins.

## INTRODUCTION

The abundance of RNA and protein molecules in a cell depends both on their rates of production and degradation. The transcription rate of RNA and the rate of degradation of proteins is determined by DNA and protein sequences, respectively (Liu et al. 2016). However, most regulatory steps that control gene expression are influenced by the sequence of the RNA itself. These processes include RNA splicing, localization, stability, and translation, all of which can be regulated by RNA-binding proteins (RBPs) that specifically recognize short RNA sequence elements (Glisovic et al. 2008).

RBPs can recognize their target sites using two mechanisms: they can form direct contacts to the RNA bases of an unfolded RNA chain, and/or recognise folded RNA-structures (reviewed in (Draper 1999; Jones et al. 2001; Mackereth and Sattler 2012)). These two recognition modes are not mutually exclusive, and the same RBP can combine both mechanisms in recognition of its target sequence. The RBPs that bind to unfolded target sequences are commonly assumed to bind to each base independently of the other bases, and their specificity is modelled by a simple position weight matrix (PWM; (Stormo 1988; Cook et al. 2011)). However, recognition of a folded RNA-sequence leads to strong positional interdependencies between different bases due to base pairing. In addition to the canonical Watson-Crick base pairs G:C and A:U, double-stranded RNA commonly contains also G:U base pairs, and can also accommodate other non-canonical base pairing configurations in specific structural contexts (Varani and McClain 2000).

It has been estimated that the human genome encodes approximately 1500 proteins that can associate with RNA (Gerstberger et al. 2014). Only some of the RBPs are thought to be sequence specific. Many RNA-binding proteins bind only a single RNA species (e.g. ribosomal proteins), or serve a structural role in ribonucleoprotein complexes or the spliceosome. As RNA can fold to complex three-dimensional structures, defining what constitutes an RBP is not simple. In this work, we have focused on identifying motifs for RBDs that bind to short sequence elements, analogously to sequence-specific DNA binding transcription factors. The number of such RBPs can be estimated based on the number of proteins containing one or more canonical RNA-binding protein domains. The total number is likely to be ~400 RBPs (Cook et al. 2011; Ray et al. 2013; Dominguez et al. 2018). The major families of RBPs contain canonical RNA-binding protein domains (RBDs) such as the RNA recognition motif (RRM), CCCH zinc finger, K homology (KH) and cold shock domain (CSD). A smaller number of proteins bind RNA using La, HEXIM, PUF, THUMP, YTH, SAM and TRIM-NHL domains (Ray et al. 2013). In addition, many “non-canonical” RBPs that do not contain any of the currently known RBDs have been reported to specifically bind to RNA (see, for example (Gerstberger et al. 2014)).

Various methods have been developed to determine the binding positions and specificities of RNA binding proteins. Methods that use crosslinking of RNA to proteins followed by immunoprecipitation and then massively parallel sequencing (CLIP-seq or HITS-CLIP, reviewed in (Darnell 2010) and PAR-CLIP (Hafner et al. 2010) can determine RNA positions bound by RBPs *in vivo*, whereas other methods such as SELEX (Tuerk and Gold 1990), RNA Bind-n-Seq (Lambert et al. 2015; Dominguez et al. 2018) and RNAcompete (Ray et al. 2009) can determine motifs bound by RBPs *in vitro*. Most high-resolution models derived to date have been determined using RNAcompete or RNA Bind-n-Seq. These methods have been used to analyze large numbers of RBPs from multiple species, including generation of models for a total of 137 human RBPs (Ray et al. 2013; Dominguez et al. 2018).

The cisBP-RNA database (Ray et al. 2013) (Build 0.6) currently lists total of 392 high-confidence RBPs in human, but contains high-resolution specificity models for only 100 of them (Ray et al. 2013). The Encyclopedia of DNA Elements (ENCODE) database that contains human RNA Bind-n-Seq data, in turn, has models for 78 RBPs (Dominguez et al. 2018). In addition, a literature curation based database RBPDB (The database of RNA-binding protein specificities) (Cook et al. 2011) contains experimental data for 133 human RBPs, but mostly contains individual target- or consensus sites, and only has high resolution models for 39 RBPs (by high resolution, we refer to models that are derived from quantitative analysis of binding to all short RNA sequences). Thus, despite the central importance of RBPs in fundamental cellular processes, the precise sequence elements bound by most RBPs remain to be determined. To address this problem, we have in this work developed high-throughput RNA SELEX (HTR-SELEX) and used it to determine binding specificities of human RNA binding proteins. Our analysis suggests that many RBPs prefer to bind structured RNA motifs, and can associate with several distinct sequences. The distribution of motif matches in the genome indicates that many RBPs have central roles in regulation of RNA metabolism and activity in cells.

## RESULTS

### Identification of RNA-binding motifs using HTR-SELEX

To identify binding specificities of human RBPs, we established a collection of canonical and non-canonical full-length RBPs and RNA binding domains, based on the presence of a canonical RBD (from cisBP-RNA database; (Ray et al. 2013)). We also included unconventional RNA-binding proteins that have been reported to bind to RNA but that lack canonical RBDs (Gerstberger et al. 2014). Full-length constructs representing 819 putative RBPs were picked from the Orfeome 3.1 and 8.1 collections (Lamesch et al. 2007). In addition, 293 constructs designed to cover all canonical RBDs within 156 human RBPs were synthesized based on Interpro defined protein domain calls from ENSEMBL v76. Most RBD constructs contained all RBDs of a given protein with 15 amino-acids of flanking sequence (see **Supplemental Table S1** for details). For some very large RBPs, constructs were also made that contained only a subset of their RBDs. Taken together our clone collection covered 942 distinct proteins (**Supplemental Table S1**). The RBPs were expressed in *E.coli* as fusion proteins with thioredoxin, incorporating an N-terminal hexahistidine and a C-terminal SBP tag (Jolma et al. 2015).

To identify RNA sequences that bind to the proteins, we subjected the proteins to HTR-SELEX (**Fig. 1A**). In HTR-SELEX, a 40 bp random DNA sequence containing a sample index and 5’ and 3’ primer binding sequences is transcribed into RNA using T7 RNA polymerase, and incubated with the individual proteins in the presence of RNase inhibitors, followed by capture of the proteins using metal-affinity resin. After washing and RNA recovery, a DNA primer is annealed to the RNA, followed by amplification of the bound sequences using a reverse-transcription polymerase chain reaction (RT-PCR) using primers that regenerate the T7 RNA polymerase promoter. The entire process is repeated up to a total of four selection cycles. The amplified DNA is then sequenced, followed by identification of motifs using the Autoseed pipeline (Nitta et al. 2015) modified to analyze only the transcribed strand (see **Methods** for details). HTR-SELEX uses a selection library with very high sequence complexity, allowing identification of long RNA binding preferences.

**Figure 1.**
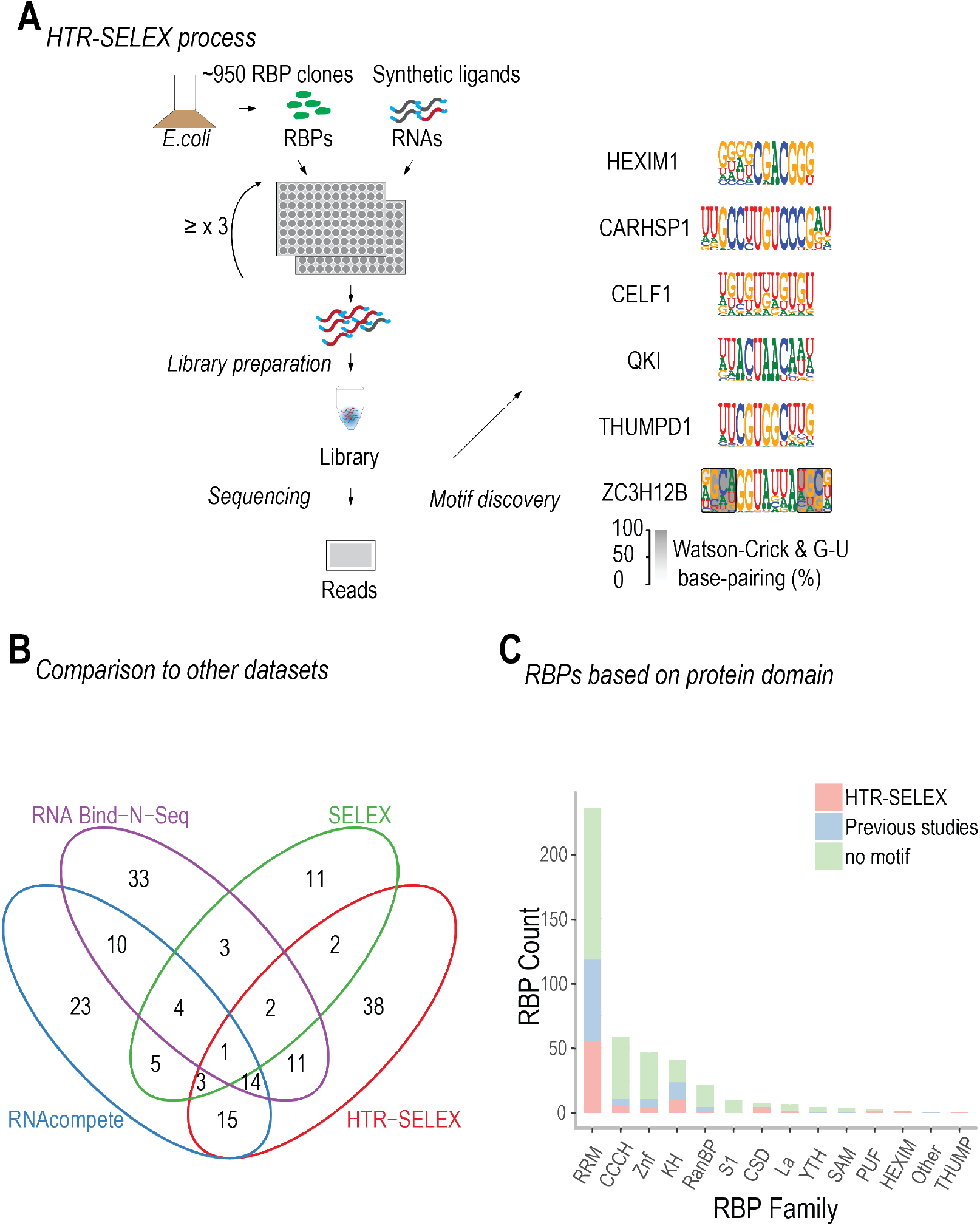
HT RNA-SELEX protocol and data-analysis. (**A**) Schematic illustration of the HTR-SELEX process. RBD or full-length RBPs expressed in *E.coli* as TRX-HIS_6_-SBP-tagged fusion proteins (top left) were purified and incubated with barcoded RNA selection ligands. RNA ligands bound by the proteins were recovered by RT-PCR, followed by *in vitro* transcription to generate the RNA for the next cycle of SELEX (left middle). The procedure was repeated at least three times and the ligands recovered from the selection cycles were subjected to Illumina sequencing (left bottom) with data analysis to generate binding specificity models (right). (**B**) Comparison of the number of RBPs with motifs derived in the present study (HTR-SELEX) with the number of RBPs for which motifs were previously derived using RNA Bind-n-Seq (RBNS) (Dominguez et al. 2018), SELEX and/or RNAcompete (cisBP-RNA version 0.6; (Ray et al. 2013)). Note that our analysis revealed motifs for 38 RBPs for which a motif was not previously known. (**C**) Distribution of RBPs with motifs classified by the structural family of their RBDs. RBPs with motifs reported in (Ray et al. 2013) and (Dominguez et al. 2018) are shown in blue, and RBPs for which motifs were not reported there but determined using HTR-SELEX in this study are in red. RBPs with no motifs are in green.

The analysis resulted in generation of 145 binding specificity models for 86 RBPs. Most of the results (66 RBPs) were replicated in a second HTR-SELEX experiment. The success rate of our experiments was ~ 22% for the canonical RBPs, whereas the fraction of the successful non-canonical RBPs was much lower (~ 1.3%; **Supplemental Table S1**). Comparison of our data with a previous dataset generated using RNAcompete (Ray et al. 2013) and RNA Bind-n-Seq (Dominguez et al. 2018) and to older data that has been compiled in the RBPDB-database (Cook et al. 2011) revealed that the specificities were generally consistent with the previous findings (**Supplemental Fig. S1** and **S2**). HTR-SELEX resulted in generation of a larger number of motifs than the previous systematic studies, and revealed the specificities of 38 RBPs whose high-resolution specificities were not previously known (**Fig. 1B**). Median coverage per RBD family was 24 % (**Fig. 1C**). Compared to the motifs from previous studies, the motifs generated with HTR-SELEX were also wider, and had a higher information content (**Supplemental Fig. S3**), most likely due to the fact that the sequences are selected from a more complex library in HTR-SELEX (see also (Yin et al. 2017)). The median width and information contents of the models were 10 bases and 10 bits, respectively. To validate the motifs, we evaluated their performance against ENCODE eCLIP data. This analysis revealed that HTR-SELEX motifs were predictive against *in vivo* data, and that their performance was overall similar to motifs generated using RNAcompete (Ray et al. 2013). The benefit of recovering longer motifs was evident in the analysis of TARDBP, whose HTR-SELEX motif clearly outperformed a shorter RNAcompete motif (**Supplemental Fig. S20**).

### Some RBPs bind to RNA as dimers

Analysis of enriched sequences revealed that 31% of RBPs (27 of 86 with an identified motif) could bind to a site containing a direct repeat of the same sequence (**Supplemental Fig. S4**, **Supplemental Tables S1 and S2**). Most of these RBPs (15 of 27) had multiple RBDs, which could bind similar sequences, as has been reported previously in the case of ZRANB2 (Loughlin et al. 2009). However, such direct repeats were also bound by RBPs having only a single RBD (12 of 27), suggesting that some RBPs could form homodimers, or interact to form a homodimer when bound to RNA (**Supplemental Table S2**). The gap between the direct repeats was generally short, with a median gap of 5 nucleotides (**Supplemental Fig. S4**). To determine whether the gap length preferences identified by HTR-SELEX were also observed in sites bound *in vivo*, we compared our data against existing *in vivo* data for four RBPs for which high quality PAR-CLIP and HITS-CLIP derived data was available from previous studies (Hafner et al. 2010; Farazi et al. 2014; Weyn-Vanhentenryck et al. 2014). We found that preferred spacing identified in HTR-SELEX was in most cases (3 out of 4) also observed in the *in vivo* data. However, the gap length distribution observed *in vivo* extended to longer gaps than that observed in HTR-SELEX (**Supplemental Fig. S5**), suggesting that such lower-affinity spacings could also have a biological role in RNA folding or function.

### Recognition of RNA structures by RBPs

Unlike double-stranded DNA, RNA folds into complex, highly sequence-dependent three dimensional structures. To analyze whether RBP binding depends on RNA secondary structure, we identified characteristic patterns of dsRNA formation by identifying correlations between all two base positions either within the motif or in its flanking regions, using a measure described in Nitta et al., (Nitta et al. 2015) that is defined by the difference between the observed count of combinations of a given set of two bases and their expected count based on a model that assumes independence of the positions (**Fig. 2A**). The vast majority of the observed deviations from the independence assumption were consistent with the formation of an RNA stem-loop structure (example in **Fig. 2B**). In addition, we identified one RBP, LARP6, that bound to multiple motifs (**Supplemental Figs. S6 and S19B**), including a predicted internal loop embedded in a double-stranded RNA stem (**Fig. 2C**). This binding specificity is consistent with the earlier observation that LARP6 binds to stem-loops with internal loops found in mRNAs encoding the collagen proteins COL1A1, COL1A2 and COL3A1 (Cai et al. 2010) (**Supplemental Fig. S6**).

**Figure 2.**
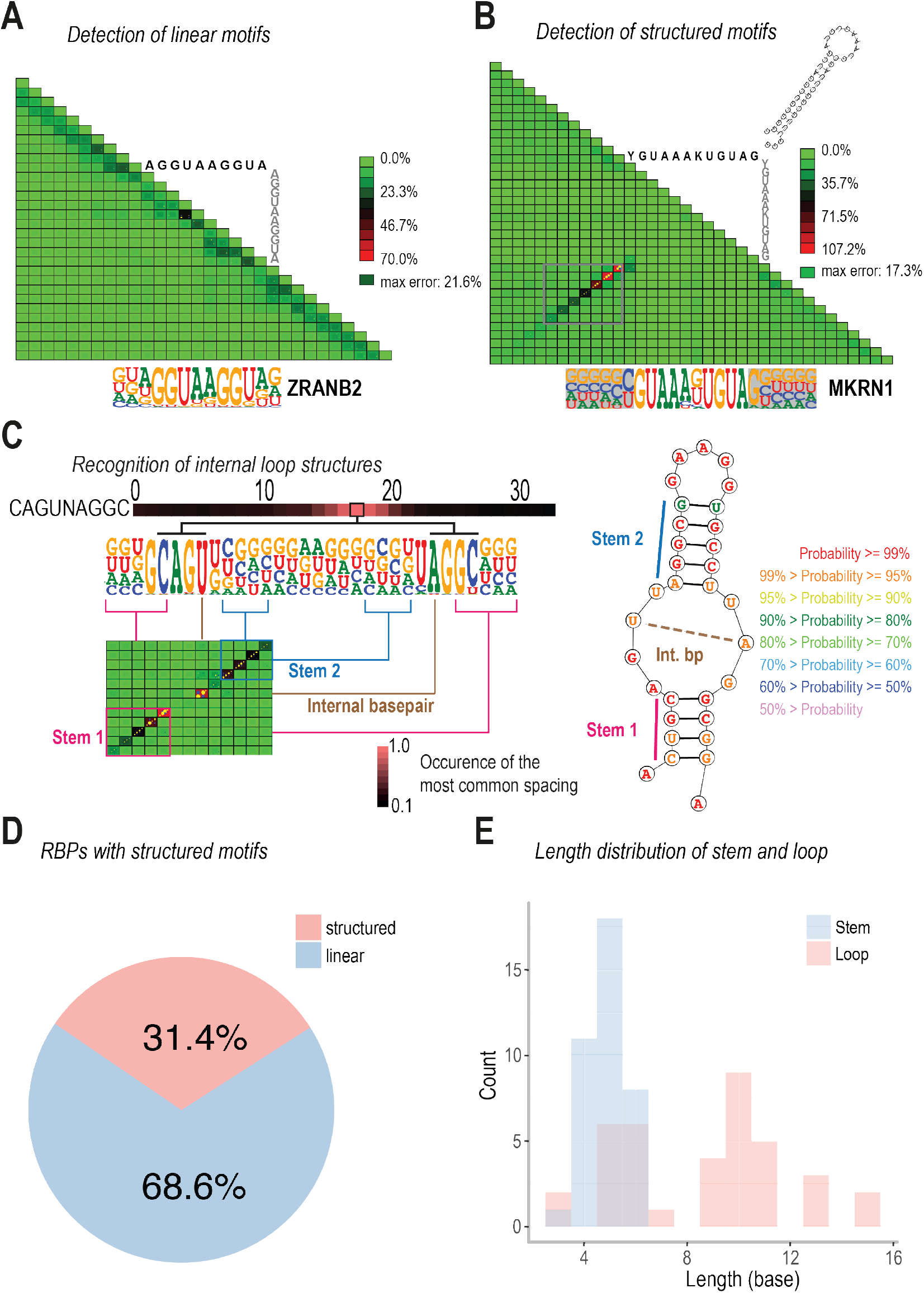
Detection of linear or structured RNA binding models. (**A**) ZRANB2 binds to a linear RNA motif. The motif of ZRANB2 and the seed used to derive it are shown below and above the triangular correlation heatmap, respectively. The heatmap illustrates deviation of the observed nucleotide distributions from those predicted by a mononucleotide model where bases are independent. (**B**) MKRN1 binds preferentially to a stem-loop. Note a diagonal series of red tiles (boxed) that indicates pairs of bases whose distribution deviates from the independence assumption. These bases are shaded in the motif below the triangle. The interdependency occurs between bases that are at the same distance from the center of the motif, consistent with formation of a stem-loop structure. Right top: A RNAfold-predicted stem-loop structure for a sequence that was highly enriched in the experiment. (**C**) LARP6 binds to a complex internal loop RNA structure. The left panel indicates the dinucleotide dependencies with the heatmap on top representing the preferred spacing length between base pairing sequences of stem 1, whereas the right panel presents a predicted structure of the bound RNA. The dashed line in the structure denotes the internal base pair. (**D**) Fraction of RBPs with linear and structured binding specificities. RBPs with at least one structured specificity are counted as structured. (**E**) Length distribution of stem and loop for the structured motifs.

In total, 69% (59 of 86) of RBPs recognized linear sequence motifs that did not appear to have a preference for a specific RNA secondary structure. The remaining 31% (27 of 86) of RBPs could bind at least one structured motif (**Fig. 2D**); this group included several known structure-specific RBPs, such as RBFOX1 (Chen et al. 2016), RC3H1, RC3H2 (Leppek et al. 2013), RBMY1E, RBMY1F, RBMY1J (Skrisovska et al. 2007) and HNRNPA1 (Chen et al. 2016; Orenstein et al. 2018). A total of 15 RBPs bound only to structured motifs, whereas 12 RBPs could bind to both structured and unstructured motifs. For example, both linear and structured motifs were detected for RBFOX proteins; binding to both types of motifs was confirmed by analysis of eCLIP data (**Supplemental Fig. S20A**).

The median length of the stem region observed in all motifs was 5 bp, and the loops were between 3 and 15 bases long, with a median length of 11 (**Fig. 2E**). Of the different RBP families, KH and HEXIM proteins only bound linear motifs, whereas proteins from RRM, CSD, Zinc finger and LA-domain families could bind to both structured and unstructured motifs (**Supplemental Fig. S7**).

To model RBP binding to stem-loop structures, we developed a simple stem-loop model (SLM; **Fig. 3; Supplemental Table S2-S4**). This model describes the loop as a position weight matrix (PWM), and the stem by a nucleotide pair model where the frequency of each combination of two bases at the paired positions is recorded. In addition, we developed two different visualizations of the model, a T-shaped motif that describes the mononucleotide distribution for the whole model, and the frequency of each set of bases at the paired positions by thickness of edges between the bases (**Fig. 3**), and a simple shaded PWM where the stem part is indicated by a gray background where the darkness of the background indicates the fraction of bases that pair with each other using Watson-Crick or G:U base pairs (**Fig. 3**). Analysis of the SLMs for each structured motif indicated that on average, the SLM increased the information content of the motifs by 4.2 bits (**Supplemental Fig. S8**). Independent secondary structure analysis performed using RNAfold indicated that as expected from the SLM, >80% of individual sequence reads for MKRN1 had more than four paired bases, compared to ~15% for the control RBP (ZRANB2) for which a structured motif was not identified (**Fig. 3**).

**Figure 3.**
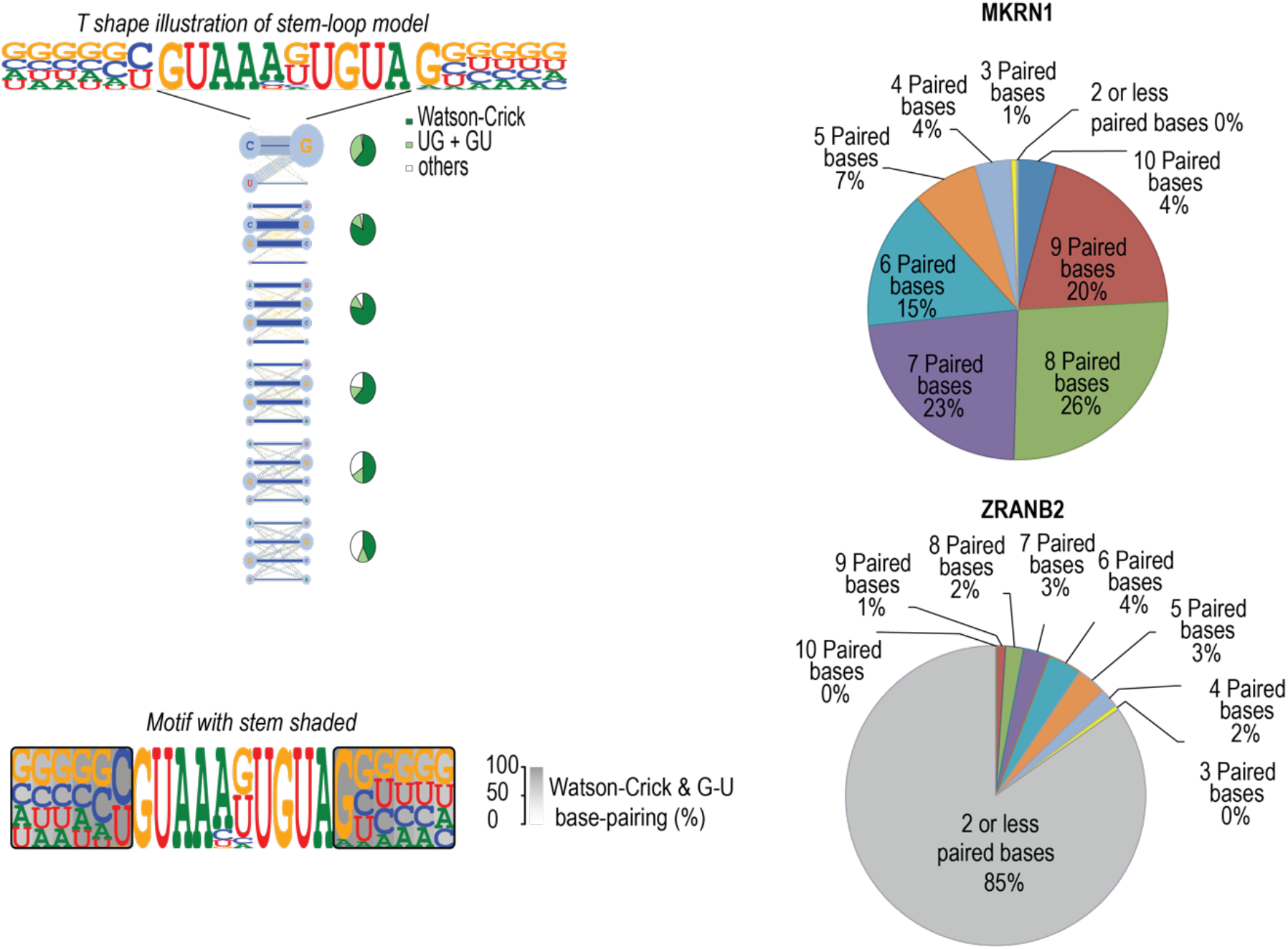
Comparison between linear PWM and stem loop (SLM) models. Left: Visualization of the stem loop models. A T-shape model (top) shows a horizontal loop and a vertical stem where the frequency of each base combination is shown. Bases are aligned so that Watson-Crick base pairs orient horizontally. Pie-charts show frequency of Watson-Crick (green) and G-U base pairs (light green) compared to other pairs (gray) that do not form canonical dsRNA base pairs at each position of the predicted stem. A linear visualization (bottom) where the base pairing frequency is indicated by the darkness of gray shading is also shown. Right: RNA secondary structure prediction analysis using RNAfold reveals that sequences flanking MKRN1 loop sequence form base pairs (top), whereas bases on the flanks of ZRANB2 matches (bottom) are mostly unpaired.

### Classification of RBP motifs

To analyze the motif collection globally, we developed PWM and SLM models for all RBPs. To compare the motifs, we determined their similarity using SSTAT (Pape et al. 2008). To simplify the analysis, PWM models were used for this comparison even for RBPs that bound to the structured motifs. We then used the dominating set method (Jolma et al. 2013) to identify a representative set of distinct motifs (**Supplemental Fig. S9**). Comparison of the motifs revealed that in general, the specificities of evolutionarily related RBPs were similar (**Fig. 4** and **Supplemental Fig. S9**). For the largest family, RRM, a total of 96 motifs were represented by 47 specificity classes, whereas the smaller families CCCH, KH, CSD, and HEXIM were represented by 9, 10, 6 and 1 classes, representing 17, 11, 7 and 2 individual motifs, respectively (**Supplemental Fig. S9**).

**Figure 4.**
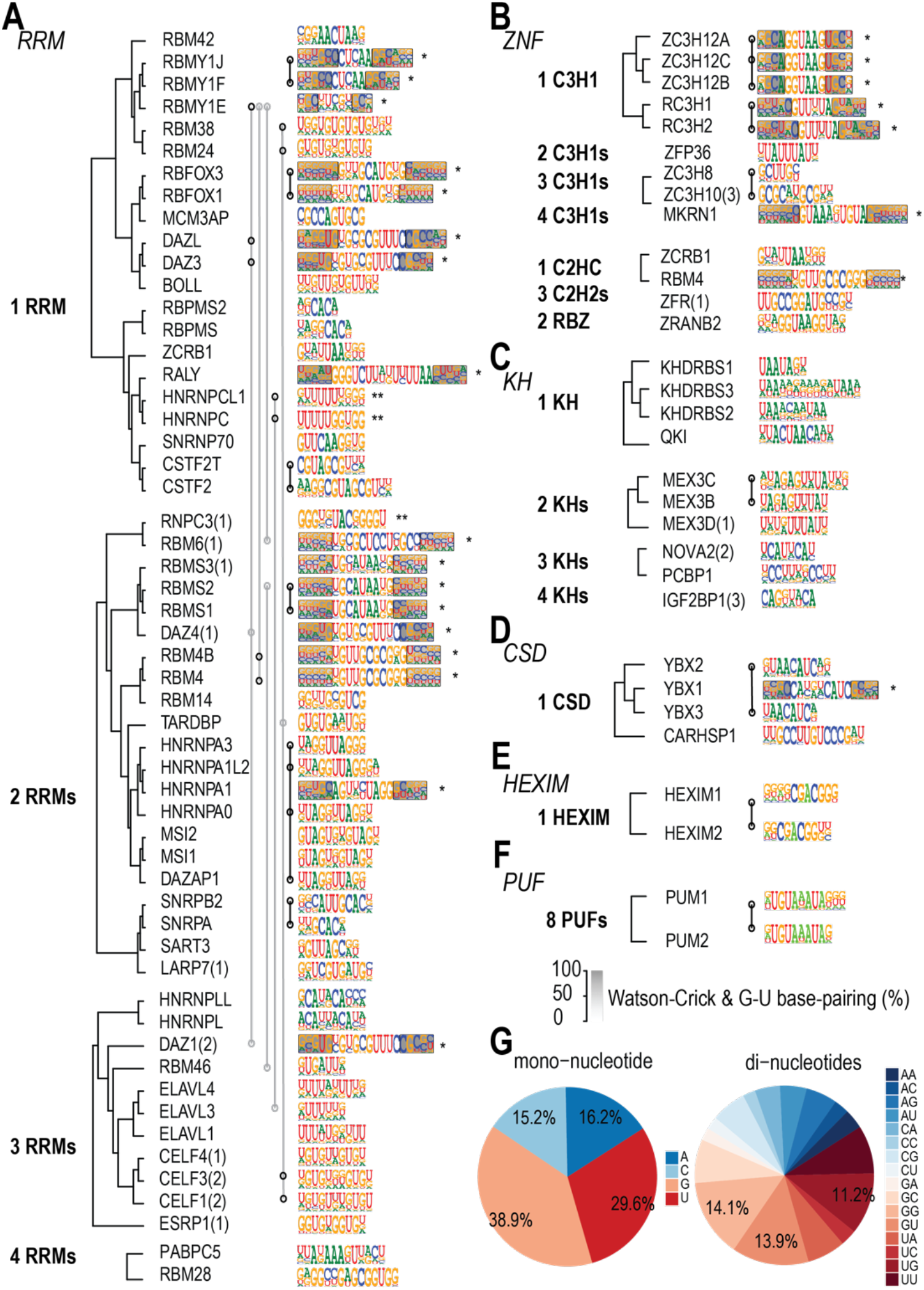
Comparison between the HTR-SELEX motifs. (**A-F**) Similar RBPs bind to similar motifs. Motifs were classified into six major categories based on structural class of the RBPs. Dendrograms are based on amino-acid alignment using PRANK. Within the RRM family, RBPs with different numbers of RRMs were grouped and aligned separately; if fewer RBDs were included in the construct used, the number of RBDs is indicated in parentheses (see also **Supplemental Table S1**). Motifs shown are the primary motif for each RBP. Asterisks indicate a stem-loop structured motif, with the gray shading showing the strength of the base pairing at the corresponding position. Two asterisks indicate that the RBP can bind to a structured secondary motif. Motifs that are similar to each other based on SSTAT analysis (covariance threshold 5 × 10^−6^) are indicated by open circles connected by lines. Only families with more than one representative HTR-SELEX motif are shown. (**G**) RBPs commonly prefer sequences with G or U nucleotides. Frequencies of all mononucleotides (left) and dinucleotides (right) across all of the RBP motifs. Note that G and U are overrepresented.

Analysis of the dinucleotide content of all motifs revealed unexpected differences in occurrence of distinct dinucleotides within the PWMs. The dinucleotides GG, GU, UG and UU were much more common than other dinucleotides (**Fig. 4G**; fold change 2.75; p < 0.00225; t-test). This suggests that G and U bases are most commonly bound by RBPs. This effect could be in part due to structural motifs, where G and U can form two different base-pairs. Furthermore, many RBPs function in splicing, and their motifs preferentially match sequences related to the G-U rich splice donor sequence A/UG:GU (**Supplemental Data S1-S4**). However, G and U enrichment cannot be explained by structure alone, as the unstructured motifs were also enriched in G and U. One possibility is that the masking of G and U bases by protein binding may assist in folding of RNA to defined structures, as G and U bases have lower specificity in base-pairing than C and A, due to the presence of the non-Watson-Crick G:U base pairs in RNA. The enrichment of G and U bases in RBP motifs was also previously reported in a different motif set discovered using a different method, RNA Bind-n-Seq (Dominguez et al. 2018). (See **Supplemental Fig. S21** for comparison with RNAcompete).

Most RBPs bound to only one motif. However, 41 RBPs could bind to multiple distinctly different motifs (**Fig. 5**). Of these, 19 had multiple RBDs that could explain the multiple specificity. However, 22 RBPs could bind to multiple motifs despite having only one RBD, indicating that individual RBPs are commonly able to bind to multiple RNA-sequences. In five cases, the differences between the primary and secondary motif could be explained by a difference in spacing between the two half-sites. In 12 cases, one of the motifs was structured, and the other linear. In addition, in eight RBPs the primary and secondary motifs represented two different structured motifs, where the loop length or the loop sequence varied (**Fig. 5**). In addition, for four RBPs, we recovered more than two different motifs. The most complex binding specificity we identified belonged to LARP6 (**Fig. 5** and **Supplemental Fig. S10**), which could bind to multiple simple linear motifs, multiple dimeric motifs, and the internal loop-structure described above.

**Figure 5.**
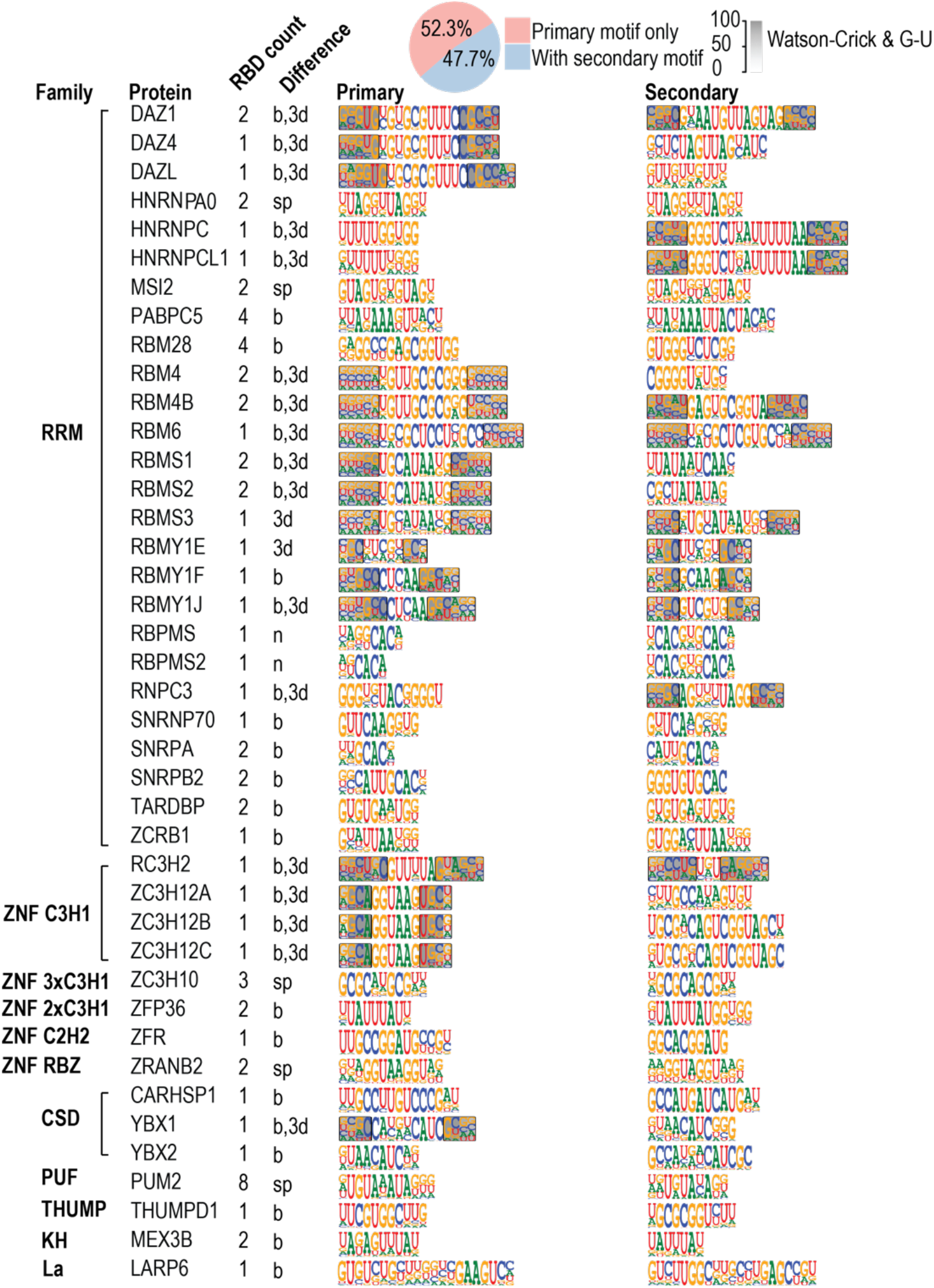
Many RBPs can recognize more than one motif. Pie chart (top) indicates fraction of RBPs that recognize more than one motif. Primary (left) and secondary (right) motifs are shown, classified according to the RBP structural family. Number next to the RBD name indicates the number of RBDs in the construct used, and the letters indicate how the two motifs are different from each other, as follows, difference in: number of half-sites (n), half-site spacing (sp), base recognition (b), and/or secondary structure (3d).

### Conservation and occurrence of motif matches

We next analyzed the enrichment of the motif occurrences in different classes of human transcripts. The normalized density of motif matches for each RBP at both strands of DNA was evaluated relative to the following features: transcription start sites (TSSs), splice donor and acceptor sites, and translational start and stop positions (see **Supplemental Fig. S11** and **Supplemental Data S1-S4** for full data). This analysis revealed that many RBP recognition motifs were enriched at splice junctions. The most enriched linear motif in splice donor sites belonged to ZRANB2, a known regulator of alternative splicing (**Fig. 6A**) (Loughlin et al. 2009). Analysis of matches to structured motifs revealed even stronger enrichment of motifs for ZC3H12A, B and C to splice donor sites (**Fig. 6A**). These results suggest a novel role for ZC3H12 proteins in regulation of splicing. The motifs for both ZRANB2 and ZC3H12 protein factors were similar but not identical to the canonical splice donor consensus sequence ag|GU[g/a]agu (**Fig. 6A**) that is recognized by the spliceosome, suggesting that these proteins may act by binding to a subset of splice donor sites.

**Figure 6.**
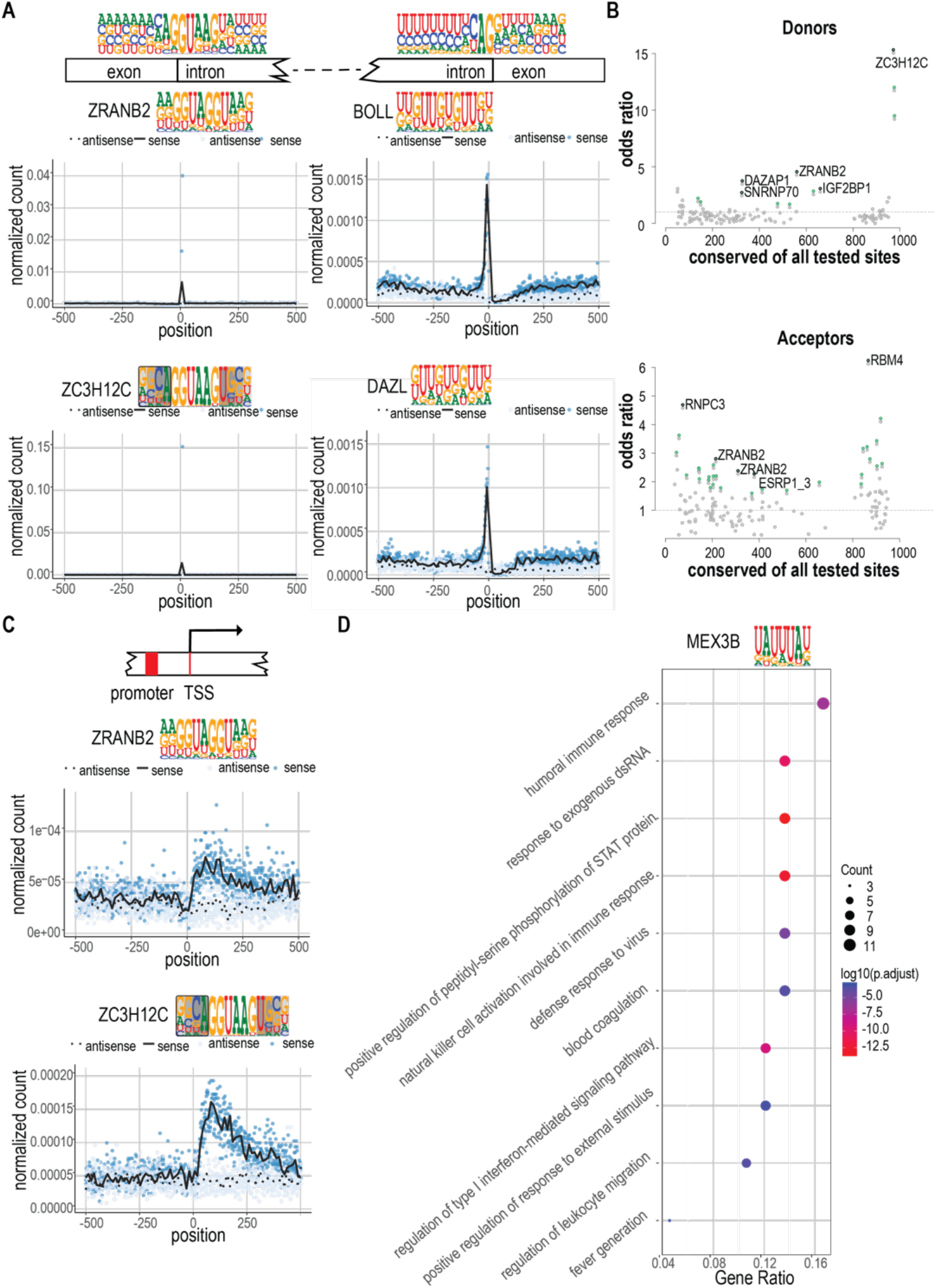
RBP motif matches are conserved and enriched in distinct sequence features and classes of transcripts. (**A**) Strong enrichment of RBP motif matches at or near the splicing donor and acceptor sites. Mononucleotide frequencies at splice donor and acceptor sites are shown on top, above the gene schematic. Left: meta-plots indicate the enrichment of ZRANB2 and ZC3H12C motif matches at splice donor sites. Right: enrichment of BOLL and DAZL at splice acceptor sites. Blue dots indicate the number of matches in the sense strand at each base position; black line indicates the locally weighted smoothing (LOESS) curve in 10 base sliding windows. Corresponding values for the anti-sense strand are shown as light blue dots and dotted black line, respectively. (**B**) The conservation of motif matches in sense vs. antisense strand. Odds ratio of preferential conservation of a match in the sense strand (y-axis) is shown as a function of the total number of conserved motif matches (x-axis; see **Methods** for details). Motifs for which conservation is significantly associated with sense strand (one-sided Fisher’s exact test) are shown in green. The five motifs with the smallest p-values are indicated in black and named. (**C**) Enrichment of ZRANB2 and ZC3H12C motif matches near transcription start sites (TSS). Note that matches are only enriched on the sense strand downstream of the TSS. (**D**) Gene Ontology enrichment of MEX3B motif matches. The top 100 genes with highest motif-matching score density were used to conduct the Gene Ontology enrichment analysis. The enriched GO terms were simplified by their similarity (cutoff=0.5). The fraction of genes and their counts in the GO categories are also shown (Gene Ratio, Count, respectively).

Analysis of splice acceptor sites also revealed that motifs for known components of the spliceosome, such as RBM28 (Damianov et al. 2006), were enriched in introns and depleted in exons (**Supplemental Data S1-S4**). Several motifs were also enriched at the splice junction, including the known regulators of splicing IGF2BP1 and ZFR (**Supplemental Data S1-S4**) (Haque et al. 2018; Huang et al. 2018). In addition, we found several motifs that mapped to the 5’ of the splice junction, including some known splicing factors such as QKI (Hayakawa-Yano et al. 2017) and ELAVL1 (Bakheet et al. 2018), and some factors such as DAZL, CELF1 and BOLL for which a role in splicing has to our knowledge not been reported (**Fig. 6A** and **Supplemental Data S1-S4**) (Rosario et al. 2017; Xia et al. 2017).

To determine whether the identified binding motifs for RBPs are biologically important, we analyzed the conservation of the motif matches in mammalian genomic sequences close to splice junctions. This analysis revealed strong conservation of several classes of motifs in the transcripts (**Fig. 6B, Supplemental Table S6**), indicating that many of the genomic sequences matching the motifs are under purifying selection.

Matches to both ZRANB2 and ZC3H12 motifs were enriched in 5’ regions of the sense-strands of known transcripts, but not on the corresponding anti-sense strands. However, no enrichment was detected in the potential transcripts that would originate from the same promoters and extend in a direction opposite to that of the mRNAs (**Fig. 6C**). These results suggest that ZRANB2 and ZC3H12 motifs could have a role in differentiating between forward and reverse strand transcripts that originate from bidirectional promoters.

We also used Gene Ontology Enrichment analysis to identify motifs that were enriched in specific types of mRNAs. This analysis revealed that many RBP motifs are specifically enriched in particular classes of transcripts. For example, we found that MEX3B motifs were enriched in genes involved in type I interferon-mediated signaling pathway (**Fig. 6D, Supplemental Table S7**).

Taken together, our analysis indicates that RBP motifs are biologically relevant, as matches to the motifs are conserved, and occur specifically in genomic features and in transcripts having specific biological roles.

### Structural analysis of ZC3H12B bound to RNA

The ability of the cytoplasmic ZC3H12 proteins to bind to splice donor-like sequences suggests that these proteins may be involved in recognition of unspliced cellular mRNA or viral transcripts in the cytoplasm, both of which would be subject to degradation. Indeed, the ZC3H12 proteins, which are conserved across metazoa, have been linked to protective responses against viral infection (Fu and Blackshear 2017; Wilamowski et al. 2018). Moreover, these proteins (and all our constructs) contain both C3H1 RBD and a PIN RNase domain, and previous studies have indicated that the ZC3H12 proteins are RNA endonucleases that rapidly degrade specific RNAs (Wilamowski et al. 2018).

To further explore our unexpected finding that these proteins are stably associated with splice donor-like sequences, we solved the structure of ZC3H12B together with a 21 base RNA sequence enriched in HTR-SELEX at 3.3Å resolution (**Fig. 7A, B**). To our surprise, we found that the RNA was bound to the PIN nuclease domain, and not to the conventional RNA-binding domain (C3H1), which was not resolved in our structure. As reported previously for ZC3H12A, ZC3H12B is a dimeric protein (Xu et al. 2012), with single Mg^2+^ ion coordinated at each active site. The dimer is held together by a relatively large contact surface (1008.2 Å^2^); however, it is predicted to exist as a monomer in solution (complex significance score CSS = 0; see also (Xu et al. 2012)). Similarly, the other contacts observed in the asymmetric unit of the crystal, including the RNA-RNA contact (877.0 Å^2^), and protein dimer-to-dimer contact (1028.1 Å^2^) appear too weak to exist in solution (CSS = 0 for both).

**Figure 7.**
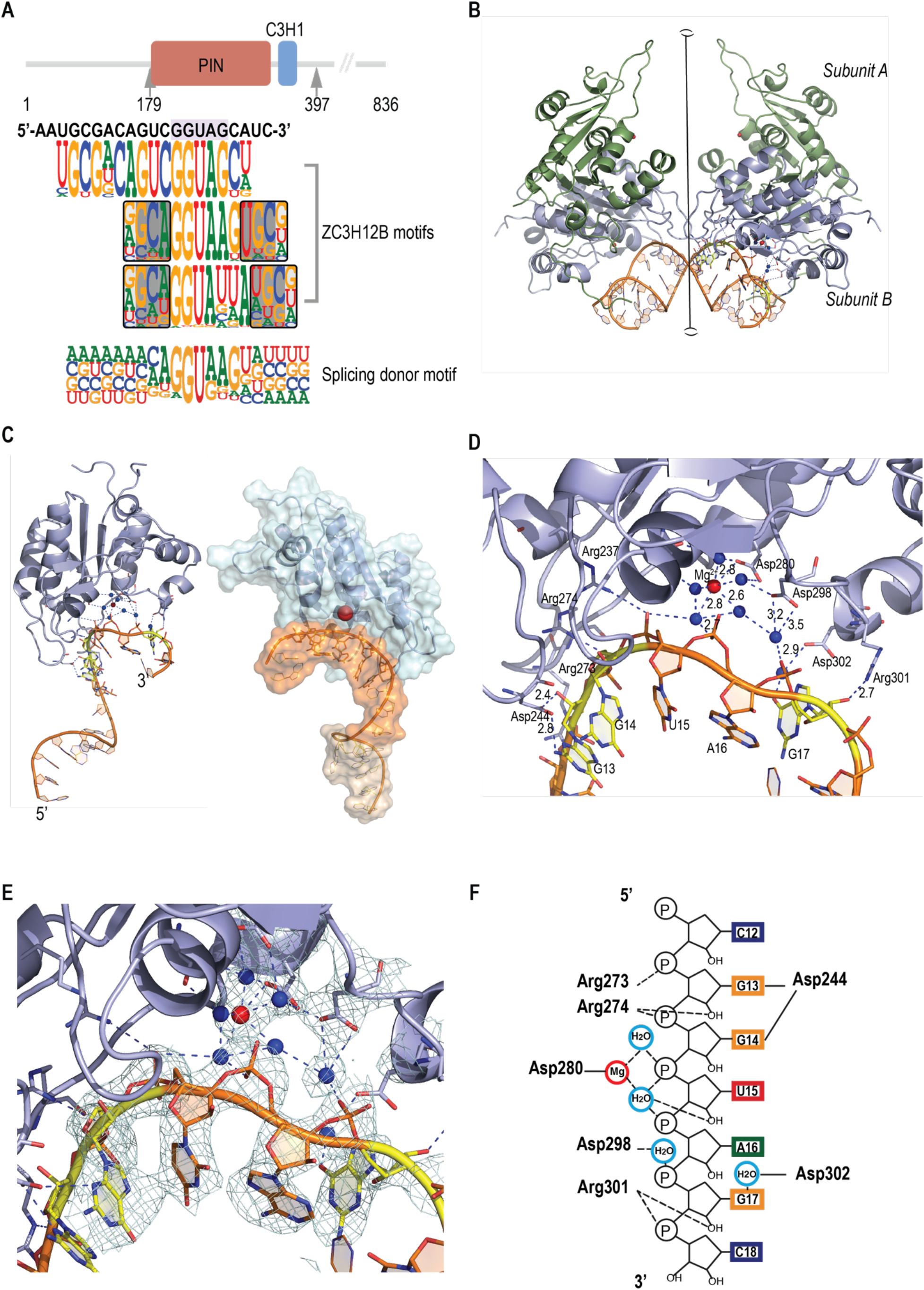
Structural basis of RNA motif recognition by ZC3H12B. (**A**) Schematic representation of the domain structure of ZC3H12B. The arrows indicate the first and the last amino acid of the construct used for crystallization, containing both the PIN domain ((Senissar et al. 2017); residues 181-350) and the known RNA-binding C3H1 Zinc finger domain ((Lai et al. 2002; Hudson et al. 2004); residues 355-380). RNA sequence used for crystallization and all ZC3H12B motifs, and the splice donor motif are shown below the cartoon. Note that all these motifs contain the sequence GGUA. (**B**) Figure shows two asymmetric units of the crystals of RNA-bound ZC3H12B. Only the PIN domain is visible in the structure. The unit belongs to P4_3_2_1_2 space group, and contains one dimer of two identical monomers presented in green (subunit A) and blue (subunit B). This dimer is similar to the dimer found in the structure of ZC3H12A (PDB: 3V33; (Xu et al. 2012)). Note that the contact between the two dimers of ZC3H12B around the 2-fold crystallographic axis (vertical line) is primarily mediated by the two RNA chains. Red and blue spheres represent Mg^2+^ ions and water molecules, respectively. For clarity, only the water molecules found in the active site are shown. Dashed lines represent hydrogen bonds (right side). The residues involved in the protein-RNA contacts are shown as ball-and stick models, and the nucleotides involved in hydrogen bonds with these residues are in yellow. Notice that only the active site of subunit B of the AB dimer is occupied by an RNA molecule. (**C**) The structure of ZC3H12B PIN domain. Left: The PIN domain is composed of a central beta-sheet surrounded by alpha-helices from both sides. The RNA molecule is bound near the Mg^2+^ ion by the -GGUAG- sequence, which is located close to the 3’ end of the co-crystallized RNA. Right: Surface model shows the shape of the active site bound by RNA (brown), with the weakly coordinated Mg^2+^ ion. Waters are omitted for clarity. Note the horseshoe-like shape of the RNA backbone at the active site (orange). (**D**) A closeup image of the RNA fragment bound to the catalytic site of ZC3H12B. Mg^2+^ ion is shown as a red sphere, the water molecules are represented as blue spheres, with dashed lines representing hydrogen bonds. Note that phosphates of U15, A16 and G17 interact with the Mg^2+^ ion via water molecules. The Mg^2+^ ion is coordinated by five water molecules also mediate contact with one of the side-chain oxygen atoms of Asp280 as well as Asp195 and Asp298 and phosphate groups of RNA. Thus, the octahedral coordination of the Mg^2+^ ion is distorted and the ion is shifted from the protein molecule towards the RNA chain, interacting with the RNA via an extensive network of hydrogen bonds. The RNA backbone is slightly bent away from the protein, suggesting that the sequence is a relatively poor substrate. The presence of only one magnesium ion and the positions of water molecules correspond to the cleavage mechanism suggested for the HIV-1 RNase H (Keck et al. 1998). (**E**) The image in D annotated with the 2Fo−Fc electron density map contoured at 1.5 σ (light green mesh). (**F**) Schematic representation of interactions between protein, the Mg^2+^ ion and RNA. Solid lines represent contacts with RNA bases, whereas hydrogen bonds to ribose and phosphates are shown as dashed lines. Nucleotide bases are presented as rectangles and colored as follows: G-yellow, A-green, U-red and C-blue. Water molecules and Mg^2+^ ion are shown as light blue and red rings, respectively.

In our structure, only one of the active sites is occupied by RNA; the protein-RNA interaction is predicted to be stable (CSS ≈ 0.6). The overall structure of the RNA-bound ZC3H12B PIN domain is highly similar to the unbound domain, and to the previously reported structure of the free PIN domain of ZC3H12A (**Supplemental Fig. S12**). The active site is relatively shallow, and the magnesium is coordinated by only one direct amino-acid contact (Asp280) together with five water molecules.

In the structure, the segment of the RNA backbone bound to the active site adopts a specific horseshoe-like shape that is highly similar to an inhibitory RNA bound to an unrelated RNase DIS3 ((Weick et al. 2018); **Supplemental Fig. S13**) in the structure of the human exosome (PDB: 6D6Q). The protein binds to five RNA bases, consistently with earlier observations suggesting that a minimum length of RNA is needed for the endonuclease activity (Lin et al. 2013). The RNA is bound mainly via interactions to the phosphate backbone and ribose oxygens; only G13 and G14 are recognized by direct hydrogen bonds between Asp244 and N3 of the guanine G13 and N3 of G14. G17, in turn, is recognized by a hydrogen bond between Arg301 and O2’ the ribose and a water-mediated hydrogen bond between Asp302 and O6 of the guanine (**Fig. 7C-F**; **Supplemental Fig. S18**). The specificity towards the central GUA trinucleotide that is common to most motifs bound by the ZC3H12 family (**Fig. 7A**) is most likely determined by an extensive water network connected to the magnesium ion, and hydrogen bonding to the symmetric molecule of RNA (G14 to U11, U15 to A9, A16 to G6; **Supplemental Fig. S14**).

The structure suggests that the RNA molecule bound to the PIN domain is a relatively poor substrate to the RNase, as although the RNA backbone is tightly bound and oriented towards the active site, the phosphate between U15 and A16 remains still relatively far from the magnesium ion.

## DISCUSSION

In this work, we have determined the RNA-binding specificities of a large collection of human RNA-binding proteins. The tested proteins included both proteins with canonical RNA binding domains and putative RBPs identified experimentally (Ray et al. 2013; Gerstberger et al. 2014). The method used for analysis involved selection of RNA ligands from a collection of random 40 nucleotide sequences. Compared to previous analyses of RNA-binding proteins, the HTR-SELEX method allows identification of structured motifs, and motifs that are relatively high in information content. The method can identify simple sequence motifs or structured RNAs, provided that their information content is less than ~40 bits. However, due to the limit on information content, and requirement of relatively high-affinity binding, the method does not generally identify highly structured RNAs that in principle could bind to almost any protein. Consistent with this, most binding models that we could identify were for proteins containing canonical RBPs.

Motifs were identified for a total of 86 RBPs. Interestingly, a large fraction of all RBPs (47%) could bind to multiple distinctly different motifs. The fraction is much higher than that observed for double-stranded DNA binding transcription factors, suggesting that sequence recognition and/or individual binding domain arrangement on single-stranded RNA can be more flexible than on dsDNA (see (Draper 1999; Jones et al. 2001; Mackereth and Sattler 2012)). Analysis of the mononucleotide content of all the models also revealed a striking bias towards recognition of G and U over C and A (see also (Dominguez et al. 2018)). This may reflect the fact that formation of RNA structures is largely based on base pairing, and that G and U are less specific in their base pairings that C and A. Thus, RBPs that mask G and U bases increase the overall specificity of RNA folding in cells.

Similar to proteins, depending on sequence, single-stranded nucleic acids may fold into complex and stable structures, or remain largely disordered. Most RBPs preferred short linear RNA motifs, suggesting that they recognize RNA motifs found in unstructured or single-stranded regions. However, approximately 31% of all RBPs preferred at least one structured motif. The vast majority of the structures that they recognized were simple stem-loops, with relatively short stems, and loops of 3-15 bases. Most of the base specificity of the motifs was found in the loop region, with only one or few positions in the stem displaying specificity beyond that caused by the paired bases. This is consistent with the structure of fully-paired double-stranded RNA where base pair edge hydrogen-bonding information is largely inaccessible in the deep and narrow major groove. In addition, we identified one RBP that bound to a more complex structure. LARP6, which has previously been shown to bind to RNA using multiple RBPs (Martino et al. 2015), recognized an internal loop structure where two base-paired regions were linked by an uneven number of unpaired bases.

Compared to TFs, which display complex dimerization patterns when bound to DNA, RBPs displayed simpler dimer spacing patterns. This is likely due to the fact that the backbone of a single-stranded nucleic acid has rotatable bonds. Thus, cooperativity between two RBDs requires that they bind to relatively closely spaced motifs.

Analysis of *in vivo* bound sequences revealed that the HTR-SELEX motifs were predictive of binding inside cells as determined by eCLIP. However, it is expected that similarly to the case of DNA-bound transcription factors, all strong motif matches will not be occupied *in vivo*. This is because binding *in vivo* will depend on competition between RBPs, their localization, and the secondary structure of the full RNAs. Analysis of the biological roles of the RBP motif matches further indicated that many motif matches were conserved, and specifically located at genomic features such as splice junctions. In particular, our analysis suggested a new role for ZC3H12, BOLL and DAZL proteins in regulating alternative splicing, and MEX3B in binding to type I interferon-regulated genes. In particular, the binding of the anti-viral cytoplasmic ZC3H12 proteins (Lin et al. 2013; Habacher and Ciosk 2017) to splice junctions may have a role in their anti-viral activity, as endogenous cytoplasmic mRNAs are depleted of splice donor sequences. As a large number of novel motifs were generated in the study, we expect that many other RBPs will have specific roles in particular biological functions.

Although we included the ZC3H12 proteins to our study because they contained the known, canonical RNA-binding domain, C3H1, our structural analysis revealed that the RNA was instead recognized specifically by the PIN domain, which has not been previously linked to sequence-specific recognition of RNA. The PIN domain active site is relatively shallow, and contains one, weakly coordinated magnesium ion. The active site was occupied by the RNA motif sequence that adopted a very specific horseshoe-like shape. The bound RNA is most likely a poor substrate for the RNase, but further experiments are needed to establish the binding affinity of and enzymatic parameters for the bound RNA species. Its binding mechanism, however, suggests that proteins containing small molecule binding pockets or active sites can bind to relatively short, structured RNA molecules that insert into the pocket. This finding indicates that it is likely that all human proteins that bind sequence-specifically to RNA motifs have not yet been annotated. In particular, several recent studies have found that many cellular enzymes bind to RNA (Hentze et al. 2018; Queiroz et al. 2019). The structure of ZC3H12B bound to RNA may thus also be important in understanding the general principles of RNA recognition by such unconventional RNA-binding proteins (Hentze et al. 2018; Queiroz et al. 2019).

Our results represent the largest single systematic study of human RNA-binding proteins to date. This class of proteins is known to have major roles in RNA metabolism, splicing and gene expression. However, the precise roles of RBPs in these biological processes are poorly understood, and in general the field has been severely understudied. The generated resource will greatly facilitate research in this important area.

## METHODS

### Clone collection, protein expression and structural analysis

Clones were either collected from the human Orfeome 3.1 and 8.1 clone libraries (full length clones) or ordered as synthetic genes from Genscript (RBP constructs). As in our previous work (Jolma et al. 2013), protein-coding synthetic genes or full length ORFs were cloned into pETG20A-SBP to create an *E.coli* expression vector that allows the RBP or RBD cDNAs to be fused N-terminally to Thioredoxin+6XHis and C-terminally to SBP-tags. Fusion proteins were then expressed in the Rosetta P3 DE LysS *E.coli* strain (Novagen) using an autoinduction protocol (Jolma et al. 2015). For protein purification and structural analysis using X-ray crystallography, see **Supplemental Methods**.

### HTR-SELEX assay

The HTR-SELEX assay was performed in 96-well plates where each well contained an RNA ligand with a distinct barcode sequence. A total of three or four cycles of the selection reaction was then performed to obtain RNA sequences that bind to the RBPs. Selection reactions were performed as follows: ~200ng of RBP was mixed on ice with ~1μg of the RNA selection ligands to yield approximate 1:5 molar ratio of protein to ligand in 20μl of Promega buffer (50 mM NaCl, 1 mM MgCl_2_, 0.5 mM Na_2_EDTA and 4% glycerol in 50 mM Tris-Cl, pH 7.5). The complexity of the initial DNA library is approximately 10^12^ DNA molecules with 40 bp random sequence (~20 molecules of each 20 bp sequence on the top strand). The upper limit of detection of sequence features of HTR-SELEX is thus around 40 bits of information content.

The reaction was incubated for 15 minutes at +37°C followed by additional 15 minutes at room temperature in 96-well plates (4-titude, USA), after which the reaction was combined with 50 μl of 1:50 diluted paramagnetic HIS-tag beads (His Mag Sepharose excel, GE-Healthcare) that had been blocked and equilibrated into the binding buffer supplemented with 0.1% Tween 20 and 0.1μg/μl of BSA (Molecular Biology Grade, NEB). Protein-RNA complexes were then incubated with the magnetic beads on a shaker for further two hours, after which the unbound ligands were separated from the bound beads through washing with a Biotek 405CW plate washer fitted with a magnetic platform. After the washes, the beads were suspended in heat elution buffer (0.5 μM RT-primer, 1 mM EDTA and 0.1% Tween20 in 10 mM Tris-Cl buffer, pH 7) and heated for 5 minutes at 70°C followed by cooling on ice to denature the proteins and anneal the reverse transcription primer to the recovered RNA library, followed by reverse transcription and PCR amplification of the ligands using primers that re-generate the T7 promoter sequences. The efficiency of the selection process was evaluated by running a qPCR reaction in parallel with the standard PCR reaction.

PCR products from RNA libraries (indexed by bar-codes) were pooled together, purified using a PCR-purification kit (Qiagen) and sequenced using Illumina HiSeq 2000 (55 bp single reads). Data was de-multiplexed, and initial data analysis performed using the Autoseed algorithm (Nitta et al. 2015) that was further adapted to RNA analysis by taking into account only the transcribed strand and designating uracil rather than thymine (for detailed description, see **Supplemental Methods**).

### Comparison of motifs and analysis of their biological function

To assess the similarity between publicly available motifs and our HTR-SELEX data, we aligned the motifs as described in (Jolma et al. 2015) (**Supplemental Fig. S1**). The alignment score for the best alignment was calculated as follows: Max (information content for PWM1 position n, information content for PWM2 position m) * (Manhattan distance between base frequencies of PWM1 position n and PWM2 position m). In regions where there was no overlap, the positions were compared to an equal frequency of all bases. The package SSTAT (Pape et al. 2008) was used to measure the similarity of the RBP PWM motifs, and the dominating set of representative motifs (see (Jolma et al. 2013)) was generated using a covariance threshold of 5 × 10^−6^.

To gain insight into the function of the RBPs, we mapped each motif to the whole human genome (hg38). We applied different strategies for the linear and the stem-loop motifs. For the linear motifs, we identified the motif matches with MOODS (Korhonen et al. 2017) with the following parameter setting: --best-hits 300000 --no-snps. For the stem-loop motifs, we implemented a novel method to score sequences against the SLMs (**Supplemental Fig. S19A**). The source code is available on GitHub: https://github.com/zhjilin/rmap.

We identified the 300,000 best scored matches in the genome, and further included any matches that had the same score as the match with the lowest score, leading to at least 300,000 matches for each motif. As the RNAs analyzed only cover 33% of the genome, this yields approx. 100,000 matches per transcriptome. The constant number of motif matches was used to make comparisons between the motifs more simple. Due to differences in biological roles of the RBPs, further analysis using distinct thresholds for particular RBPs is expected to be more sensitive and more suitable for identifying particular biological features.

The matches were then intersected with the annotated features from the ENSEMBL database (hg38, version 91), including the splicing donor (DONOR), splicing acceptor (ACCEPTOR), the translation start codon (STARTcodon), the translation stop codon (STOPcodon) and the transcription starting site (TSS). The above features were filtered in order to remove short introns (<50bp), and features with non-intact or non-canonical start codon or stop codon. The filtered features were further extended 1kb both upstream and downstream in order to place the feature in the centre of all the intervals. The motif matches overlapping the features were counted using BEDTOOLS (version 2.15.0) and normalized by the total number of genomic matches for the corresponding motif. For analysis of conservation of motif matches, mutual information analysis, and Gene Ontology enrichment, see **Supplemental Methods**.

## DATA ACCESS

All next generation sequencing data have been deposited to European Nucleotide Archive (ENA) under Accession PRJEB25907. The diffraction data and the model of the ZC3H12B:RNA complex are deposited with the Protein Data Bank under accession code 6SJD. All computer programs and scripts used are either published or available upon request. Requests for materials should be addressed to J.T.

## ACKNOWLEDGEMENTS

We thank Drs. Minna Taipale and Bernhard Schmierer for the critical review of the manuscript as well as Sandra Augsten, Lijuan Hu and Anna Zetterlund for the technical assistance. The work was supported by a travel and project grant support (Es.M., L.J.M.) from the Mayo Clinic - Karolinska Institutet collaboration partnership as well as the Knut and Alice Wallenberg Foundation (KAW 2013.0088) and the Swedish Research Council (Postdoctoral grant, 2016-00158).

## AUTHOR CONTRIBUTIONS

J.T., A.J and L.J.M. designed the experiments; A.J., Es.M. and Y.Y. performed the SELEX experiments; A.J., J.Z., J.T., K.L, T.K., T.R.H., Q.M. and F.Z. analyzed the data; Ek.M. and G.B. solved the structure; J.T., A.J. and J.Z. wrote the manuscript.

