## Supplemental Materials for "Binding specificities of human RNA binding proteins towards structured and linear RNA sequences"

### Supplemental Methods

#### Protein production

All constructs described in **Supplemental Table S1** were expressed in Rosetta P3 DE LysS *E.coli* using the autoinduction protocol described in (Jolma et al. 2015). Proteins were then purified using HIS-tag based IMAC purification. Protein production was assessed in parallel by 96-well SDS-PAGE (ePage, Invitrogen; see **Supplemental Fig. S16**). The success rate of protein production was dependent on the size of the proteins, with most small RBDs expressing well in *E.coli*. Significantly lower yield of protein was observed for full-length proteins larger than 50 kDa. All proteins were subjected to HTR-SELEX, regardless of protein level expressed. For interim storage, glycerol was added to a final concentration of 10%. Samples were split to single-use aliquots with approximately 200 ng RBP in a 5μl volume and frozen at -80°C. Expression and purification of the RNA-binding domain fragment of human ZC3H12B including PIN (residues 179-354) and Zn-finger (residues 355-397) domains was performed as described in (Savitsky et al. 2010).

#### Protein-RNA complex crystallization

The fragment of RNA composed from 21 ribonucleotides used in crystallization trials was obtained from IDT. The RNA (sequence: 5'-A*AUGCGACAGUCGGUAGCAUC-3') was protected from non-specific RNases by phosphorothioation of the 5' end (bond containing sulphur indicated by *). The purified and concentrated protein was first mixed with a solution of RNA at a molar ratio 1:1.2 and after one hour on ice it was subjected to the crystallization trials with several crystallization screens from different vendors. The first crystals of 0.04 mm size were obtained in Nuc-Pro HTS screen from Jena Bioscience and they diffracted to 6Å only. The conditions for bigger crystals were optimized in house. Crystals of 0.1 mm size were grown in sitting drops from solution containing 50 mM Sodium cacodylate buffer (pH 6.5), 80 mM $MgCl_2$ and 5% MPD (2-Methyl-2,4-pentanediol). The lifetime of the crystals was short (not more than 48 hours after the crystallization was set), suggesting that the RNA species was slowly hydrolyzed, possibly by the ZC3H12B RNase itself. The single data set was collected at the beamline P13 at PETRA- III (EMBL, Hamburg, Germany) at 100 K. Data were processed and analyzed with the autoPROC toolbox (Vonrhein et al. 2011) including the STARANISO routine due to the high anisotropy (Tickle 2017). The datasets were indexed and integrated by XDS and scaled together with XSCALE. Statistics of data collection are presented in **Supplemental Table S5**.

### Structure determination and refinement

The structure was solved by molecular replacement using the program Phaser (McCoy et al. 2007) as implemented in CCP4 (Winn et al. 2011) with the structure of ZC3H12A PIN-domain (pdb:3V33) as a search model. The density of the part of RNA was clear near the active site and the molecule was built manually using COOT (Emsley et al. 2010). The rigid body refinement with REFMAC5 was followed by restrain refinement with REFMAC5, as implemented in CCP4 (Winn et al. 2011) and Phenix.refine (Afonine et al. 2012). The manual rebuilding of the model was performed using COOT. The refinement statistics are presented in Supplemental Table S5. The first five amino acids from N-termini and the Zn-finger part from C-termini were found disordered and were not built in the maps. The four last nucleotides of the 3'- end were not visible in the maps as well, thus, were not built to the map also. The protein model was validated using COOT and MOLPROBITY (Chen et al. 2010). The structural figures were prepared using PyMOL(TM) Molecular Graphics System (Version 1.8.6.0, Schrödinger, LLC).

### Selection library generation

To produce a library of RNA sequences for selection (selection ligands), we first constructed dsDNA templates by combining three oligonucleotides together in a three cycle PCR reaction (Phusion, NEB). For information about the barcoded ligand design, see **Supplemental Table S1**. The ligand design was similar to that used in our previous work analyzing TF binding specificities in dsDNA (Jolma et al. 2013) except for the addition of a T7 RNA polymerase promoter in the constant flanking regions of the ligand. RNA was expressed from the DNA-templates using T7 *in vitro* transcription (Ampliscribe T7 High Yield Transcription Kit, *Epicentre* or Megascript-kit *Ambion*) according to manufacturer's instructions, after which the DNA-template was digested using RNAse-free DNAse I (Epicentre) or the TURBO-DNAse supplied with the Megascript-kit. All RNA-production steps included RiboGuard RNAse-inhibitor (Epicentre).

Two different approaches were used to facilitate the folding of RNA molecules. In the protocol used in experiments where the batch identifier starts with letters "EM", RNA-ligands were heated to +70°C followed by gradual, slow cooling to allow the RNA to fold into minimal energy structures, whereas in batches "AAG" and "AAH" RNA transcription was not followed by such folding protocol. The rationale was that spontaneous co-transcriptional RNA-folding may better reflect folded RNA structures in the *in vivo* context. In almost all of the cases where the same RBPs were tested with both of the protocols the results were highly similar.

**Motif generation**

The motifs were generated based on Autoseed; Autoseed identifies gapped and ungapped kmers that represent local maximal counts relative to similar sequences within their Huddinge neighborhood (Nitta et al. 2015). It then generates a draft motif using each such kmer as a seed. This procedure makes each generated PWM motif distinctly different from any other motif derived from the same data, by ensuring that the count for each seed is higher than that of any subsequence that is shifted by one base, or within a Hamming distance of one from the seed (see **Supplemental Fig. S19B**, and Supplementary Figure 1 of Nitta et al., 2015). It is important to note that the resulting motifs are not generated from a single set of aligned sequences, and that therefore the count for the base representing the consensus sequence is constant, whereas the total counts in each column vary (Jolma et al. 2013).

This initial set of motifs is then refined manually to identify the final seeds (**Supplemental Table S2**). The manual curation process was necessary to remove artefacts due to selection bottlenecks (low complexity libraries), partial motifs that included constant linker sequences (displayed a strong positional bias on the ligand), and motifs that were recovered from a large number of experiments; the motifs recovered from many experiments were removed because they represent common "aptamer" motifs that are enriched by the HTR-SELEX process itself, for example due to residual presence of *E.coli* derived RNA-binding proteins, or binding of folded "aptamer" RNAs to the TRX fusion partner, selection beads or plasticware (**Supplemental Fig. S19C**). To assess initial data, we compared the deduced motifs to known motifs, to replicate experiments (same experiment run again or separate experiment using full-length and RBD clones) and experiments performed with paralogous proteins. We also note that each cycle of HTR-SELEX independently enriches motifs over the input cycle, providing further evidence of reproducibility. Individual results that were not supported by replicate or prior experimental data were deemed inconclusive and were not included in the final dataset. Draft models were manually curated (by AJ, JT, QM, TRH) to remove unsuccessful experiments and artefacts due to bottlenecks and aptamer selection (see above), and final models were generated using the seeds indicated in **Supplemental Table S2**.

Autoseed detected more than one seed for many RBPs. Up to four seeds were used to generate a maximum of two unstructured and two structured motifs. Of these, the motif with largest number of seed matches using the multinomial setting indicated on **Supplemental Table S2** was designated the primary motif. The motif with the second largest number of matches was designated the secondary motif. The counts of the motifs represent the prevalence of the corresponding motifs in the sequence pool (**Supplemental Table S2**). Only these primary and secondary motifs were included in further analyses. Such additional motifs are shown for LARP6 in **Supplemental Fig. S10**.

To find RBPs that bind to dimeric motifs, we visually examined the PWMs to find direct repeat pattern of three or more base positions, with or without a gap between them (see **Supplemental Table S2**). The presence of such repetitive pattern could be either due to dimeric binding, or the presence of two RBDs that bind to similar sequences in the same protein.

To identify structured motifs, we visually investigated the correlation diagrams for each seed to find motifs that displayed the diagonal pattern evident in **Fig. 2B**. The plots display effect size and maximal sampling error, and show the deviation of nucleotide pair distribution from what is expected from the distribution of the individual nucleotides. For each structured motif, SLM models (**Supplemental Table S3**) were built from sequences matching the indicated seeds; a multinomial 2 setting was used to prevent the paired bases from influencing each other. Specifically, when the number of occurrences of each pair of bases was counted at the base-paired positions, neither of the paired bases was used to identify the sequences that were analyzed. The SLMs were visualized either as the T-shaped logo (**Fig. 3**) or as a PWM type logo where the bases that constitute the stem were shaded based on the total fraction of A:U, G:C and G:U base pairs.

To control for potential secondary structure bias introduced by the constant linker regions, we used the program RNAfold (Lorenz et al. 2011) to fold a set of full ligands, containing 40 bp random sequences flanked by the linkers. This analysis revealed that the constant linkers did not impose a stereotypic secondary structure on the random sequence (**Supplemental Fig S15**), indicating that the random sequences can adopt many secondary structures that are known to be important for RBP binding, such as stem-loops and internal loops, even in the context of the flanking constant linkers.

For analysis of RNA structure in **Fig. 2** and **Supplemental Fig. S6**, sequences matching the regular expression NNNNCAGU[17N]AGGCNNN or sequences of the three human collagen gene transcripts (From 5' untranslated and the beginning the coding sequence, the start codon is marked with bold typeface: COL1A1 -CCACAAAGAGUCUACAUGUCUAGGGUCUAG-AC**AUG**UUCAGCUUUGUGG; COL1A2- CACAAGGAGUCUGCAUGUCUAAGUGCUAGA-C**AUG**CUCAGCUUUGUG and COL3A1 - CCACAAAGAGUCUACAUGGGUC**AUG**UUCAG-CUUUGUGG) were analyzed using "RNAstructure" software (Mathews 2014) through the web-interface in:http://rna.urmc.rochester.edu/RNAstructureWeb/Servers/Fold/Fold.html using default settings. All structures are based on the program's minimum energy structure prediction. For analysis in **Fig. 3**, we extracted all sequences that matched the binding sequences of MKRN1 and ZRANB2 (GUAAAKUGUAG and NNNGGUAAGGUNN, respectively; N denotes a weakly specified base) flanked with ten bases on both sides from the cycle four of HTR-SELEX. Subsequently, we predicted their secondary structures using the program RNAfold (Vienna RNA package; (Lorenz et al. 2011)) followed by counting the predicted secondary structure at each base position in the best reported model for each sequence. For both RBPs, the most common secondary structure for the bases within the defined part of the consensus (GUAAAKUGUAG and GGUAAGGU) was the fully single stranded state (82% and 30% of all predicted structures, respectively). To estimate the secondary structure at the flanks, the number of paired bases formed between the two flanks were identified for each sequence. Fraction of sequences with specific number of paired bases are shown in **Fig. 3**.

### GO analysis and *in vivo* enrichment of the motifs

To determine whether RBPs with similar RBDs recognize and bind to similar targets, we compared the sequences of the RBDs and their motifs. First, the RBPs were classified based on the type and number of RBDs. For each class, we then extracted the amino-acid sequence of the RBPs starting from the first amino acid of the first RBD and ending at the last amino acid of the last RBD. We also confirmed the annotation of the RBDs by querying each amino acid sequence against that SMART database, and annotated the exact coordinates of the domains through the web-tools: http://smart.embl-heidelberg.de and http://smart.embl-heidelberg.de/smart/batch.pl. Sequence similarities and trees were built using PRANK (Loytynoja and Goldman 2005) (parameters: -d, -o, -showtree). The structure of the tree representing the similarity of the domain sequence was visualized using R (version 3.3.1).

For identification of classes of transcripts that are enriched in motif matches for each RBP, we extracted the top 100 transcripts according to the score density of each RBP motif. These 100 transcripts were compared to the whole transcriptome to conduct the GO enrichment analysis for each motif using the R package ClusterProfiler (version 3.0.5).

To analyze conservation of motif matches, sites recognized by each motif were searched from both strands of 100 bp windows centered at the features of interest (acceptor, donor sites) using the MOODS program (version 1.0.2.1). For each motif and feature type, 1000 highest affinity sites were selected for further analysis regardless of the matching strand. Whether the evolutionary conservation of the high affinity sites was explained by the motifs was tested using program SiPhy (version 0.5, task 16, seedMinScore 0) and multiz100way multiple alignments of 99 vertebrate species to human (downloaded from UCSC genome browser, version hg38). A site was marked as being conserved according to the motif if its SiPhy score was positive meaning that the aligned bases at the site were better explained by the motif than by a neutral evolutionary model (hg38.phastCons100way.mod obtained from UCSC genome browser). Two motifs (see Supplemental Table S2) were excluded from the analysis because the number of high affinity sites that could be evaluated by SiPhy was too small. The hypothesis that the motif sites in the sense strand were more likely to be conserved than sites in the antisense strand was tested against the null hypothesis that there was no association between site strand and conservation using Fisher's exact test (one-sided). The P values given by the tests for individual motifs were corrected for multiple testing using Holm's method. We note here that evidence obtained using this method establishes that the sequence under the motif matches is under purifying selection (not evolving according to the used neutral model), and is more conserved than the sequence under reverse complement matches. However, it can still be that the match sequences have another function, which can be either related (binding to a related protein) or unrelated (binding to a different class of regulator, e.g. spliceosome) to the biological mechanism of interest (RNA binding by the RBP protein).

To assess the utility of the produced motifs in predicting *in vivo* target sites (**Supplemental Fig. S20)**, they were used to predict bound sequences in eCLIP from the ENCODE portal (Davis et al. 2018). To compare HTR-SELEX with established methods, peaks from eCLIP experiments (see **Supplemental Table S8** for the accession numbers and details of the used datasets) were downloaded for proteins which had both an HTR-SELEX motif and an available RNAcompete motif on the CISBP-RNA database (Ray et al. 2013). RBFOX1 motifs were used in prediction of RBFOX2 eCLIP peaks as previous analysis has indicated that the proteins have identical RRMs (Chen et al. 2016). All peaks were extended by 20 bases upstream to account for RBP binding at the 5' end of the peak. A control set was created by taking length-matched sequences 300 bases upstream of each extended peak. The eCLIP peaks and control sets were scanned using the HTR-SELEX and RNAcompete motifs, and the max score per sequence was taken. The ability of the motif to discriminate between the two sets was evaluated by calculating the area under the ROC curve (AUROC).

The preference of RBFOX1 for binding to a hairpin loop structure was determined by first folding the eCLIP peaks and control sequences using RNAfold with the "-p" option to determine the centroid structure (Lorenz et al. 2011). Before folding, 50 flanking bases were added to both ends of each sequence to provide greater context for defining the structure and were removed after folding. Occurrences of the sequence "GCAUG" in peak and control sets were counted within hairpin loops and in other structural contexts.

### Calculation of mutual information

The global pattern of motifs across the features tested was analyzed by calculating the mutual information (MI) between 3-mer distributions at two non-overlapping positions of the aligned RNA sequences. MI can be used for such analysis, because if a binding event contacts two continuous or spaced 3-bp wide positions of the sequences at the same time, the 3-mer distributions at these two positions will be correlated. Such biased joint distribution can then be detected as an increase in MI between the positions.

Specifically, MI between two non-overlapping positions (pos1, pos2) was estimated using the observed frequencies of a 3-mer pair (3+3-mer), and of its constituent 3-mers at both positions:

$$MI(pos1, pos2) = \sum P(3{+}3\text{-}mer) \log_2 \frac{P(3{+}3\text{-}mer)}{P_{pos1}(3\text{-}mer) P_{pos2}(3\text{-}mer)}$$

where $P(3{+}3\text{-}mer)$ is the observed probability of the 3-mer pair (i.e. gapped or ungapped 6 mer). $P_{pos1}(3\text{-}mer)$ and $P_{pos2}(3\text{-}mer)$, respectively, are the marginal probabilities of the constitutive 3-mers at position 1 and position 2. The sum is over all possible 3-mer pairs.

To focus on RBPs that specifically bind to a few closely related sequences, such as RBPs with well-defined motifs, it is possible to filter out most background non-specific bindings (e.g., selection on the shape of RNA backbone) by restricting the MI calculation, to consider only the most enriched

3-mer pairs for each two non-overlapping positions. Such enriched 3-mer pair based mutual information (E-MI) is calculated by summing MI over top-10 most enriched 3-mer pairs.

$$E\text{-}MI(pos1, pos2) = \sum_{top\ 3+3\text{-}mers} P(3+3\text{-}mer) \log \frac{P(3+3\text{-}mer)}{P_{pos1}(3\text{-}mer) P_{pos2}(3\text{-}mer)}$$

### Supplemental Figures

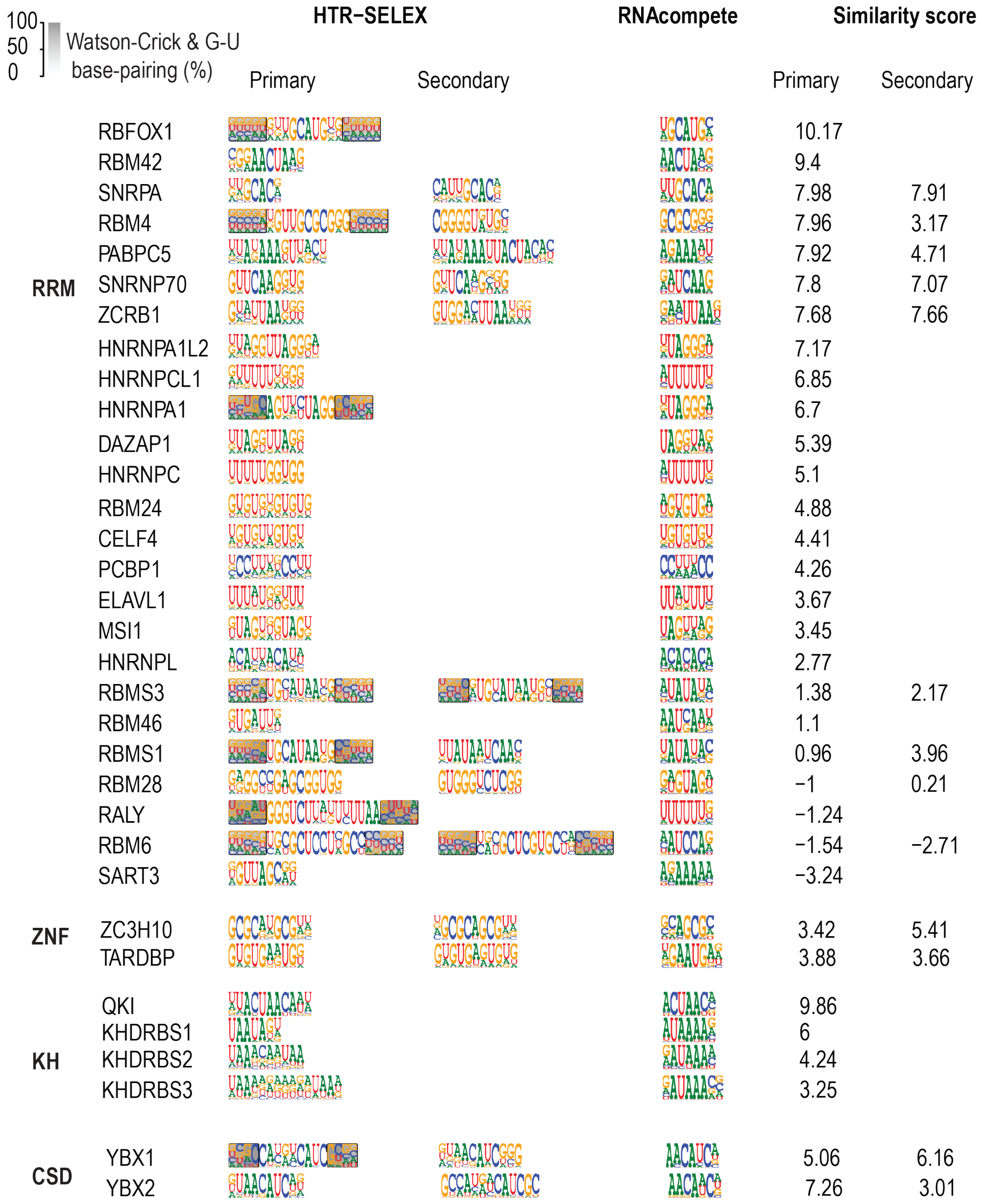

**Supplemental Figure S1. The similarity of motifs between HTR-SELEX and RNAcompete.** Comparison of HTR-SELEX and RNAcompete generated motifs for all 33 proteins for which motifs were obtained using both methods. Comparison includes both primary and secondary HTR-SELEX motifs. Motifs are organized according to protein structural family, each of which is further ordered by motif alignment score (see **Methods**). Higher score indicates higher similarity between motifs.

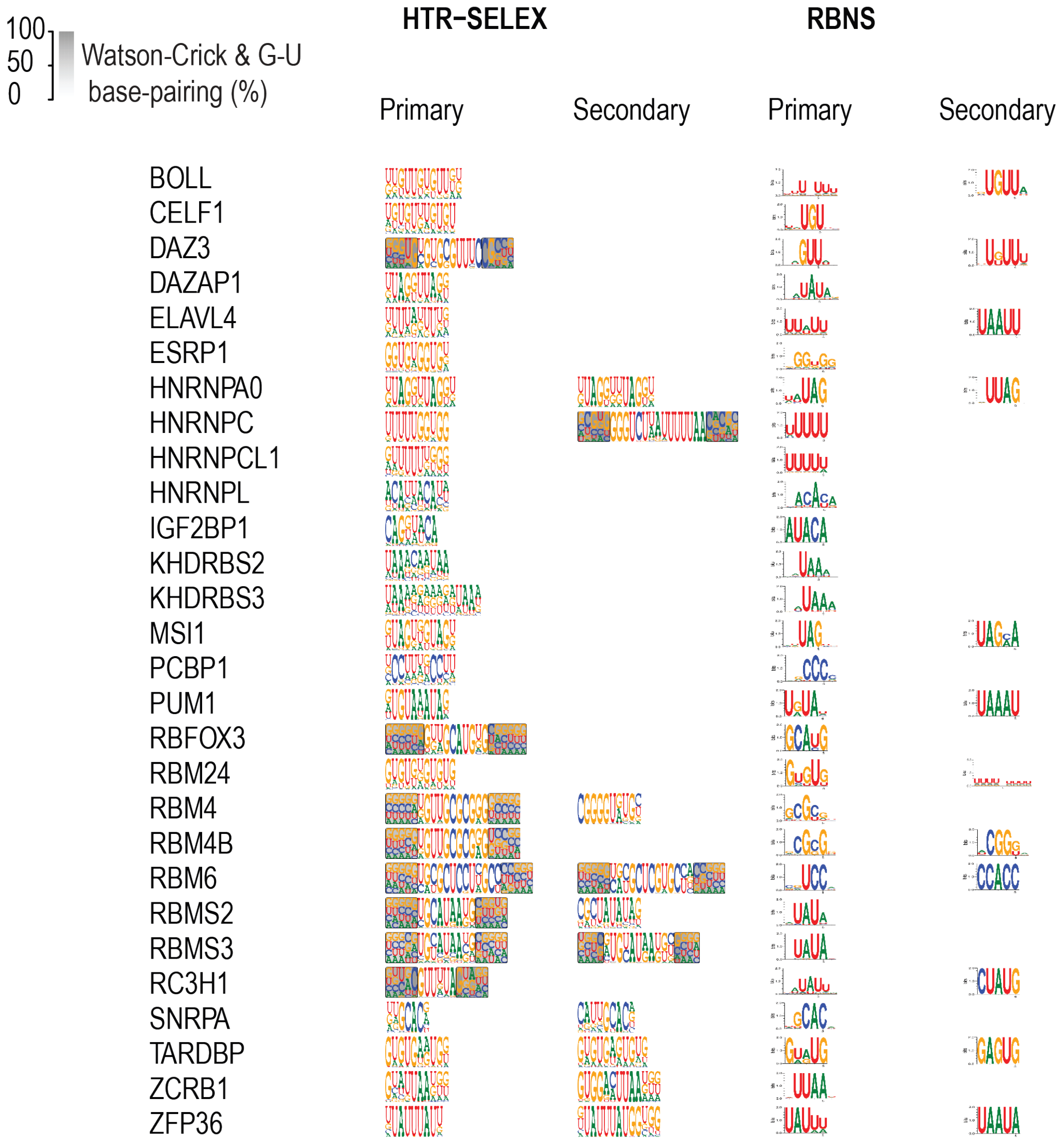

**Supplemental Figure S2. Motif comparison between HTR-SELEX and RNA Bind-n-Seq.** Comparison of HTR-SELEX and RNA Bind-n-Seq - generated motifs for all 28 proteins for which motifs were obtained using both methods. The primary and secondary motifs recovered from both experiments are presented; for RNA Bind-n-Seq, the primary motifs were determined by the most abundant 5-mers in the selected kmer population.

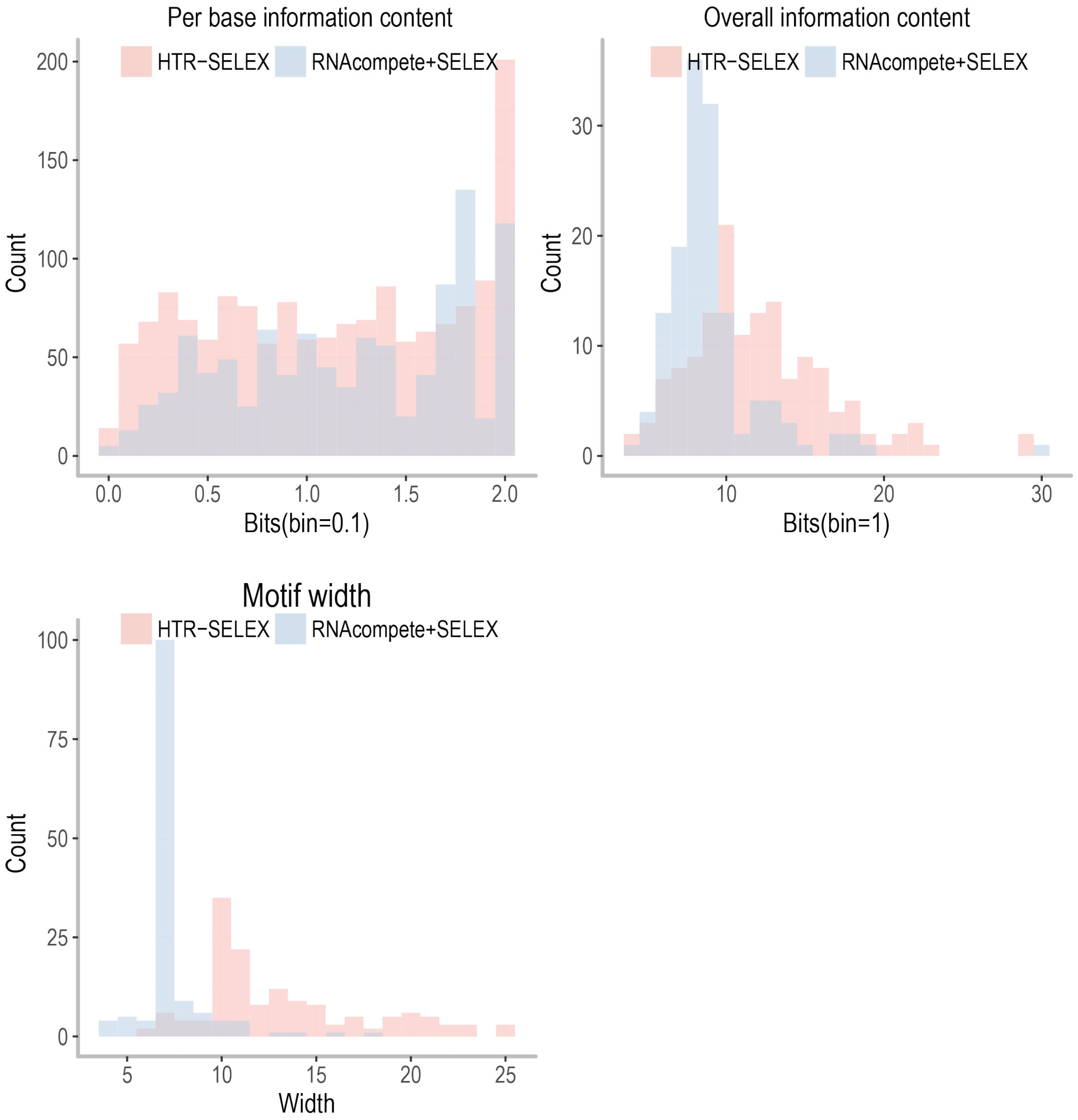

**Supplemental Figure S3. Higher information content and wider width distribution of HTR-SELEX motifs.**

The available PWMs generated by RNAcompete and SELEX were collected from the CISBP-RNA database for comparison. The per base information was calculated for every individual position in the PWM. The overall information content of each motif is the sum of all positions in the PWM. The width of each motif was generated by counting the number of position in the corresponding PWM.

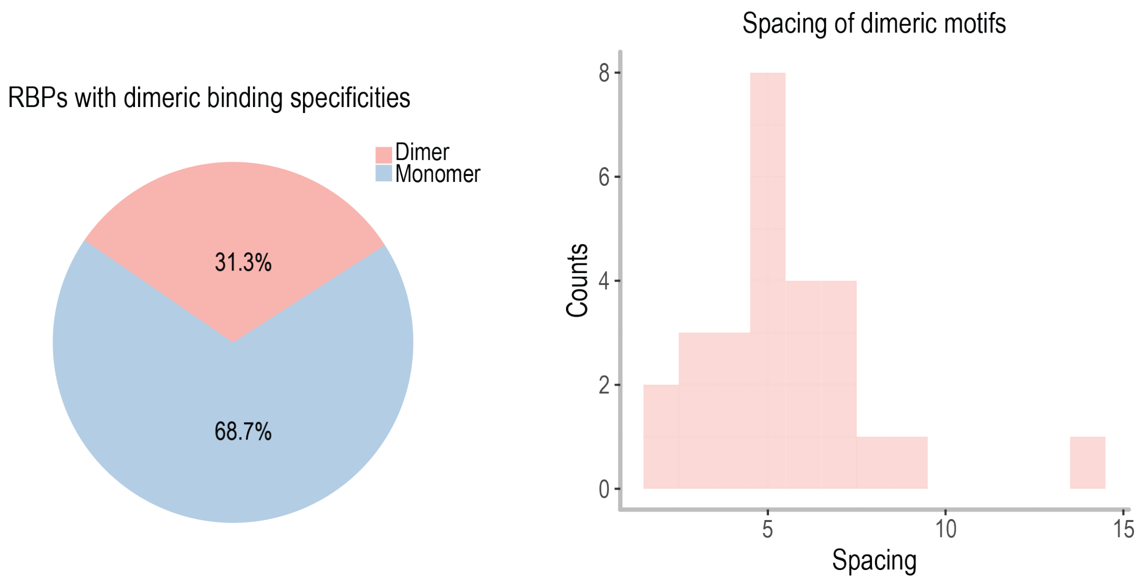

**Supplemental Figure S4. RBPs with multimeric binding sites.** About one third of RBPs (31.3%, left) bind to the sequence as homodimers where two identical half-sites are separated by a spacing sequence. The distribution of spacing preference of all RBPs is shown (right). The discontinuous distribution of the spacing length is due to the small sample size.

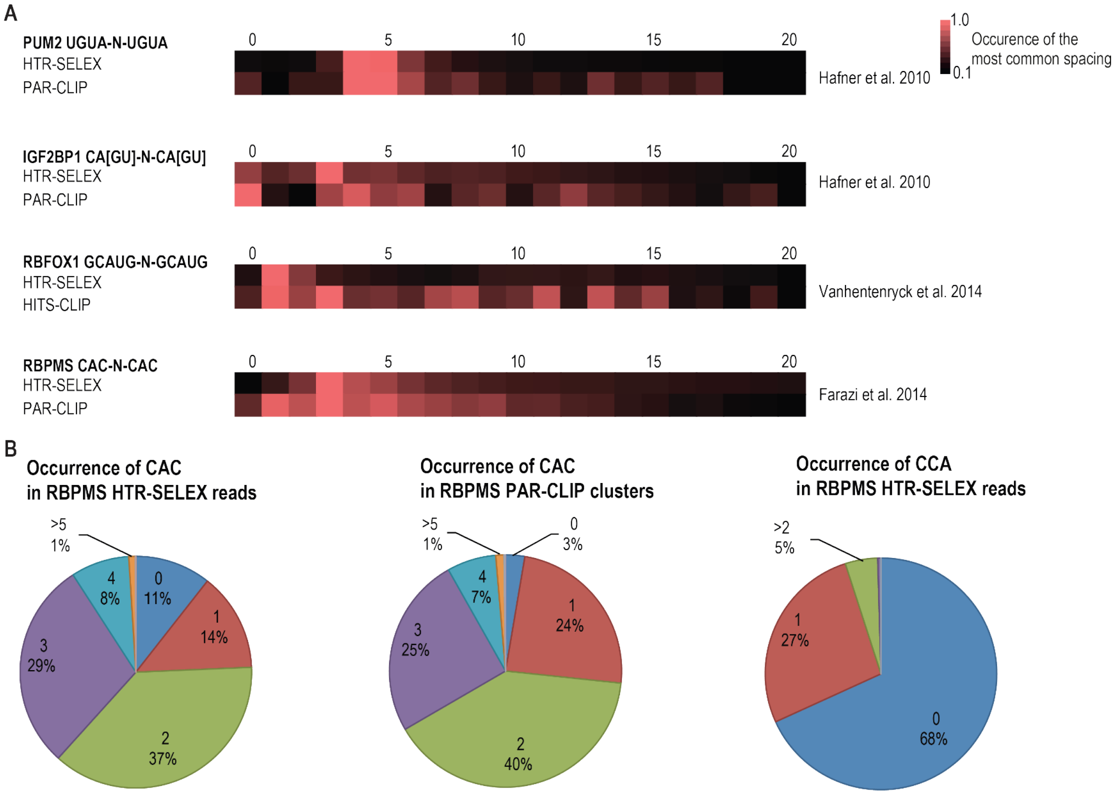

**Supplemental Figure S5. Spacing preferences between dimeric binding sites are consistent in different assays.**

**(A)** For four RBPs, the same seeds were used in different assays to detect the spacing preferences. The heatmaps represent the spacing information extracted from HTR-SELEX, PAR-CLIP and HITS-CLIP. The results are consistent between HTR-SELEX (top row) and PAR-CLIP or HITS-CLIP (bottom row). (**B**) Pie charts show the percentage of reads containing the indicated number of matches to CAC sequence in RBPMS target sequences as determined by HTR-SELEX (left) or PAR-CLIP (middle). Occurrence of the CCA-sequence that is not recognised by RBPMS but has the same base content is also shown as an example of randomly expected incidence (right).

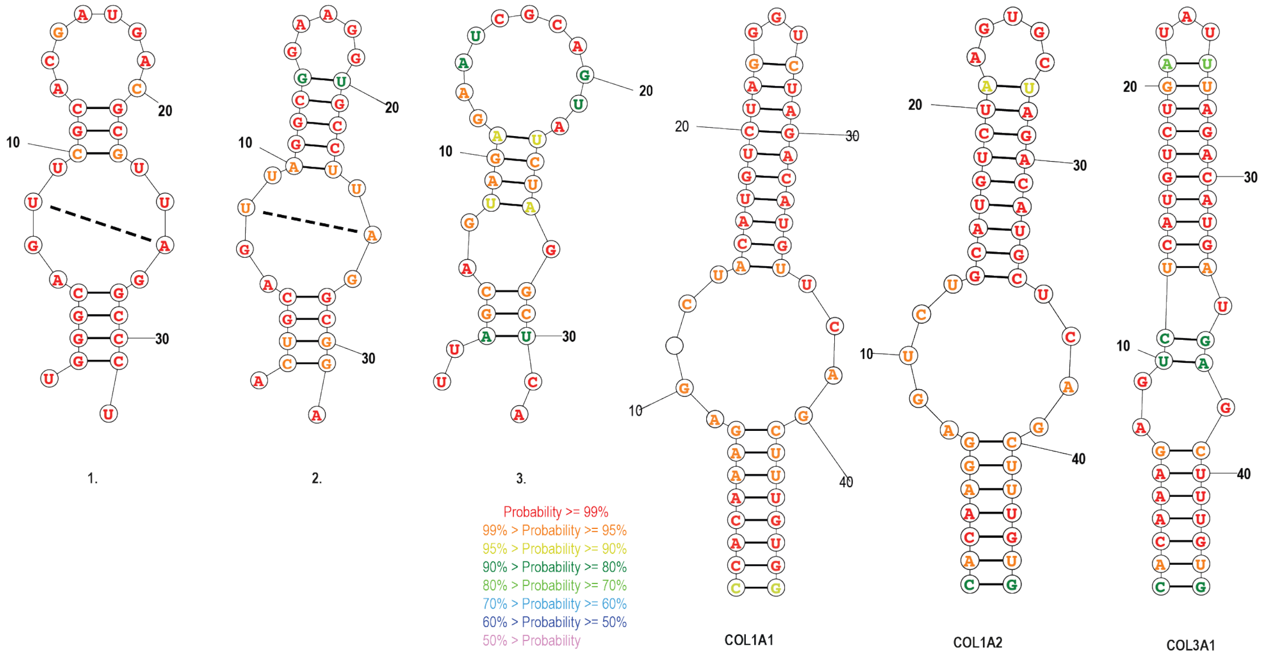

**Supplemental Figure S6. Known binding motifs of LARP6.**

The left three structures were generated using the sequences enriched in HTR-SELEX. The right three structures illustrate the predicted structures of known collagen RNA sequences. The dash-line indicates the internal base pair. The number labels the position of the base in the RNA sequence.

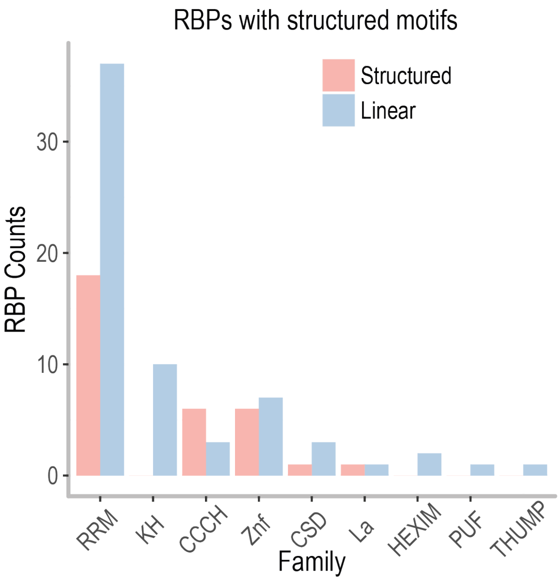

**Supplemental Figure S7. RBP families with and without structural specificity.**

The count of RBPs recognizing structured and unstructured binding motifs in each protein structure family.

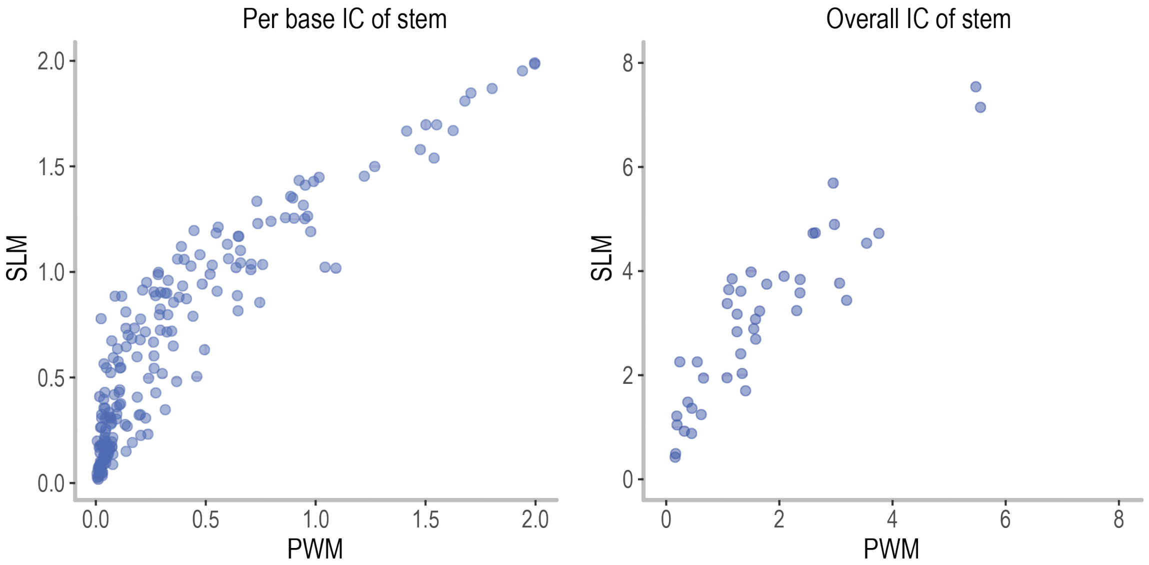

**Supplemental Figure S8. Information content correlation between the SLM and the mono-nucleotide PWM.**

Left. Information content correlation per base. Right. Overall information content correlation. In general, the SLM yielded higher per base information content due to the base pairing in the stem.

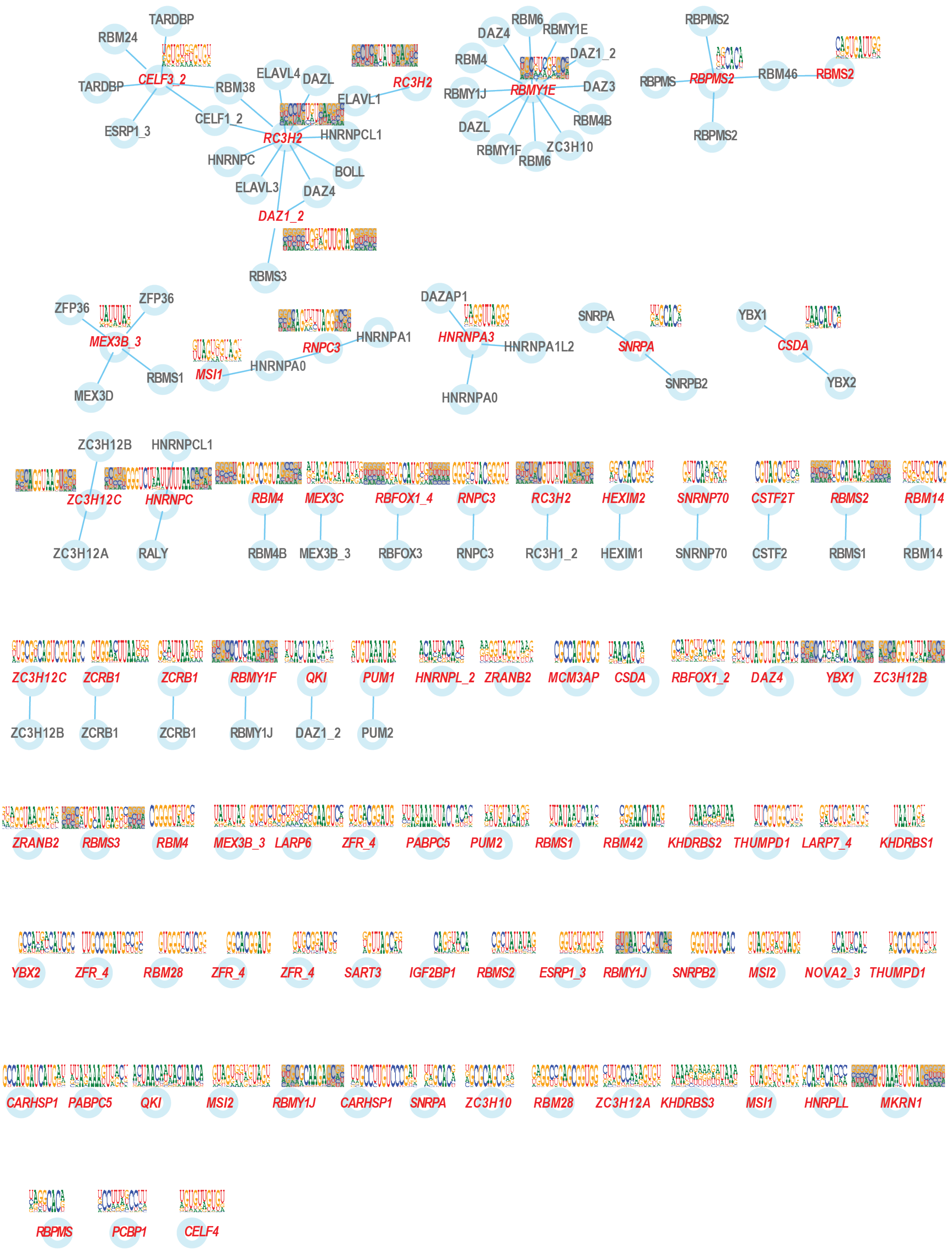

**Supplemental Figure S9. Dominating set of HTR-SELEX motifs.**

Cystoscope (Version 3.2.1) was used to visualize the dominating set on top of the relationship map between motifs with a cutoff of 5e-6 for similarity, calculated by SSTAT (see the method part). Motifs in the dominating set are labeled in red.

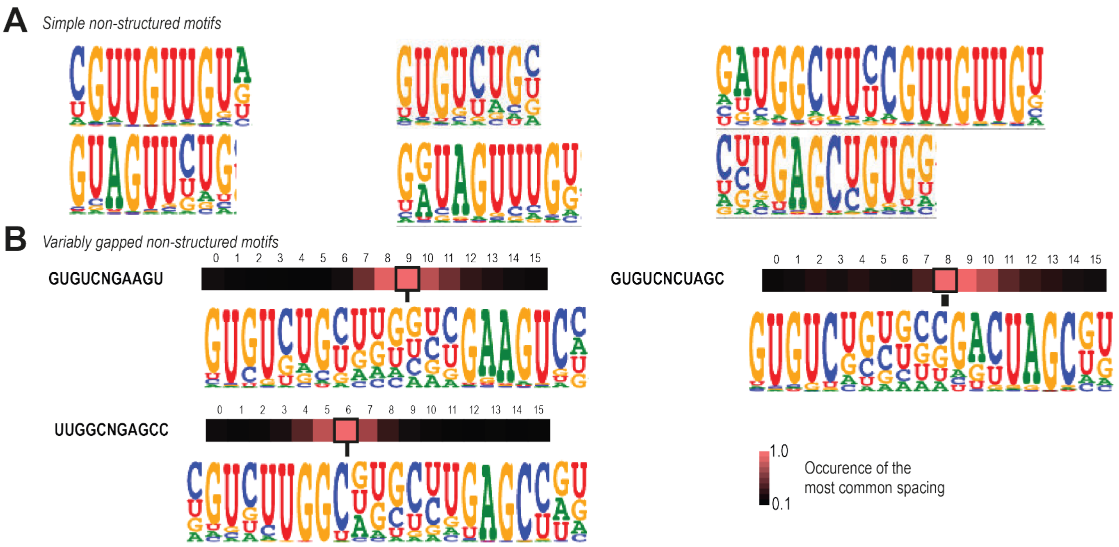

**Supplemental Figure S10. Various binding specificities detected for LARP6.**

LARP6 is able to recognise and bind to distinct sequences through different strategies besides binding to the internal loop structure. (**A**) Short and long linear motifs (**B**) unstructured motifs with gaps. The heatmap shows the preference of spacing between two half sites.

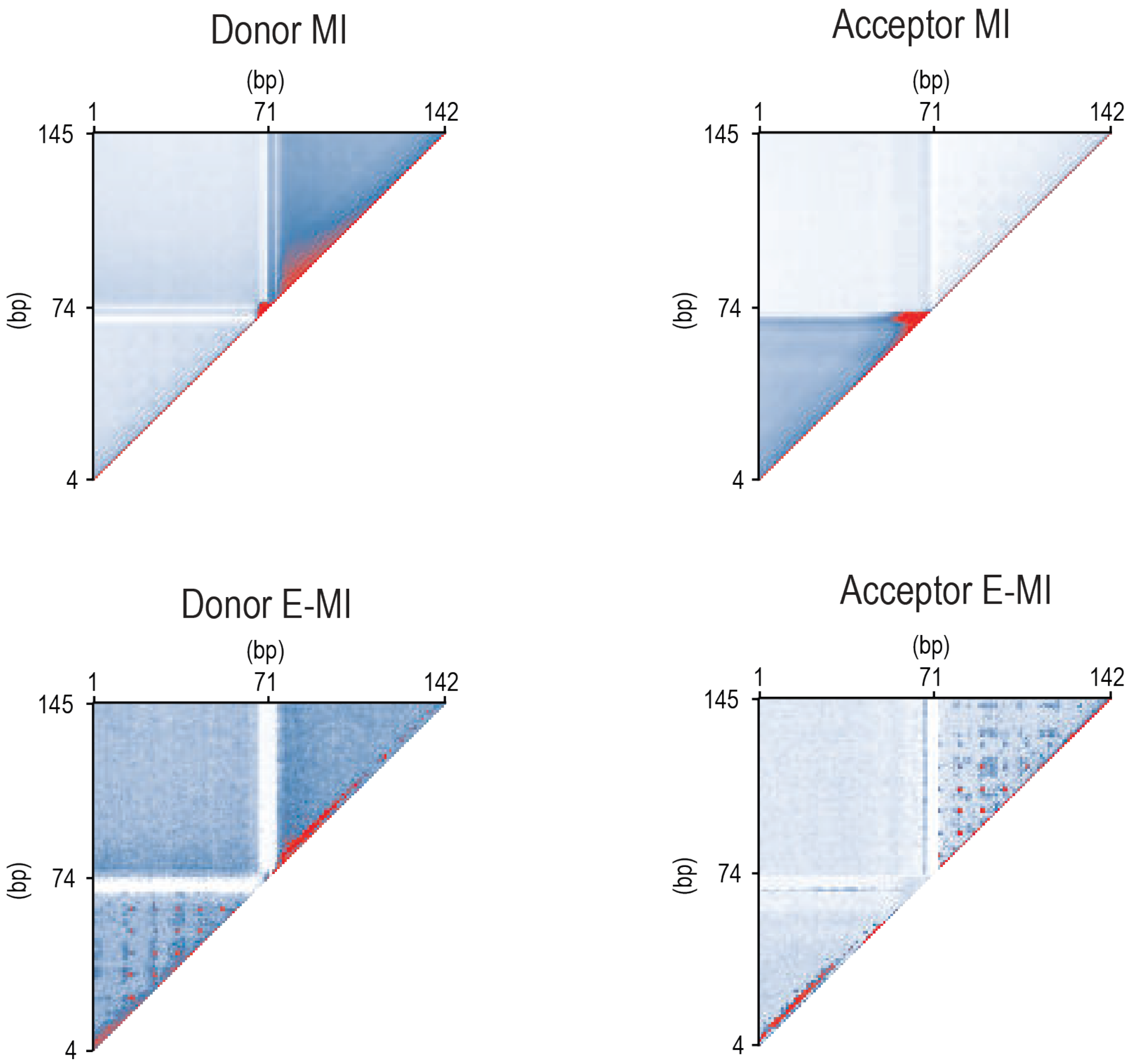

**Supplemental Figure S11. The mutual information (MI) meta-plots around the splicing donor and acceptor sites.**

The splice donor and acceptor sites are placed in the centre of the 147nts sequence. The detected signals close to the donor and acceptor sites are shown in red. The enriched 3-mer pair based mutual information (E-MI) are also shown.

A

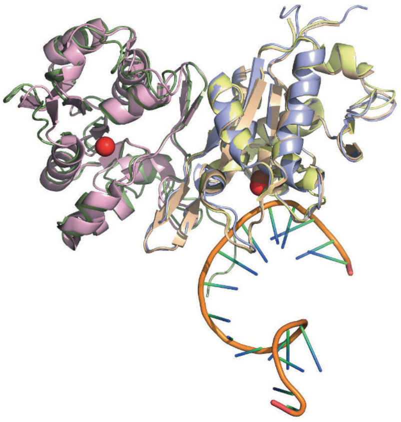

B

```
                    ß1   α1                      α2
ZC3H12B   -------------SLEDEIDNSDNLRPVVIDGSNVAMSHGNKEEFSCRGIQLAVDWFLDKG 225
ZC3H12A   GGGTPKAPNLEPPLPEEEKEGSDLRPVVIDGSNVAMSHGNKEVFSCRGILLAVNWFLERG 171

            ß2     α3           α4          ß3                  α5
ZC3H12B   HKDITVFVPAWRKEQSRPDAPITDQDILRKLEKEKILVFTPSRRVQGRRVVCYDDRFIVK 285
ZC3H12A   HTDITVFVPSWRKEQPRPDVPITDQHILRELEKKKILVFTPSRRVGGKRVVCYDDRFIVK 231

           α5    ß4        α6          α7          ß5    ß6           α8
ZC3H12B   LAFDSDGIIVSNDNYRDLQVEKPEWKKFIEERLLMYSFVNDKFMPPDDPLGRHGPSLENF 345
ZC3H12A   LAYESDGIVVSNDTYRDLQGERQEWKRFIEERLLMYSFVNDKFMPPDDPLGRHGPSLDNF 291

ZC3H12B   LRKRPIVPEHKKQPCPYGKKCTYGHKCKYYHPERANQPQRSVADELRISAK  397
ZC3H12A   LRKKPLTLEHRKQPCPYGRKCTYGIKCRFFHPERPSCPQRSVA--------  334
```

**Supplemental Figure S12. The comparison of ZC3H12B with the homologous protein ZC3H12A.**

**(A)** Superimposition of homodimer ZC3H12B (colored in green and blue) with the respective dimer from ZC3H12A NCD-ZF (colored in pink and yellow, respectively) and ZC3H12A NCD monomer containing a $Mg^{2+}$ ion (colored beige, the $Mg^{2+}$ ion is presented as brown sphere). Overall the structures are very similar (rmsd = 0.456Å and 0.439 Å, respectively). The difference is observed in the loop areas and in the slightly shifted position of the $Mg^{2+}$ ion. **(B)** The sequence alignment of ZC3H12B and ZC3H12A performed with Clustal Omega (Sievers et al. 2011). Sequence numbering is presented in the right side of the sequences. The secondary structure elements correspond to ZC3H12B are named on the top and highlighted in yellow (α-helixes) and blue (β-strands). The residues involved in the interactions with RNA are shown in bold violet and highlighted by green boxes. Two aspartate residues (Asp280 and Asp298) involved the Mg-ion coordination are colored red. The sequence corresponds to Zn-finger is highlighted grey for both proteins.

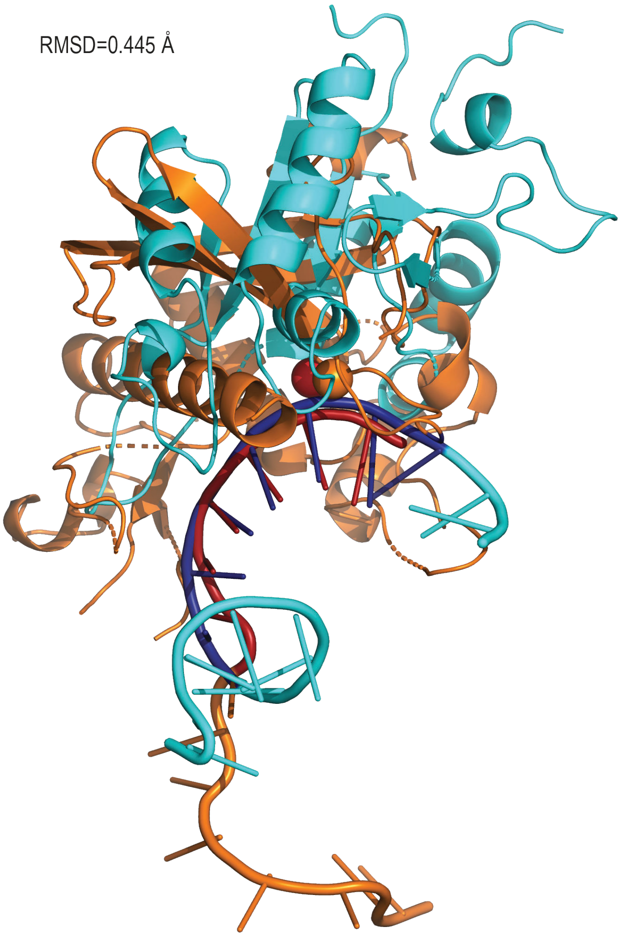

**Supplemental Figure S13. Comparison of ZC3H12B:RNA and DIS3:RNA structures.**

Superimposition of ZC3H12B:RNA (cyan) structure with a structure of the DIS3:RNA (orange) complex from the human exosome bound to an inhibitory nucleic acid. Note that the protein structures of ZC3H12B and DIS3 are completely different, whereas the overall horseshoe-like structure of the bound RNA is very similar near the active site. The RNA segments folded into the horseshoe-shape in complexes with ZC3H12B and DIS3 are indicated dark blue and dark brown, respectively.

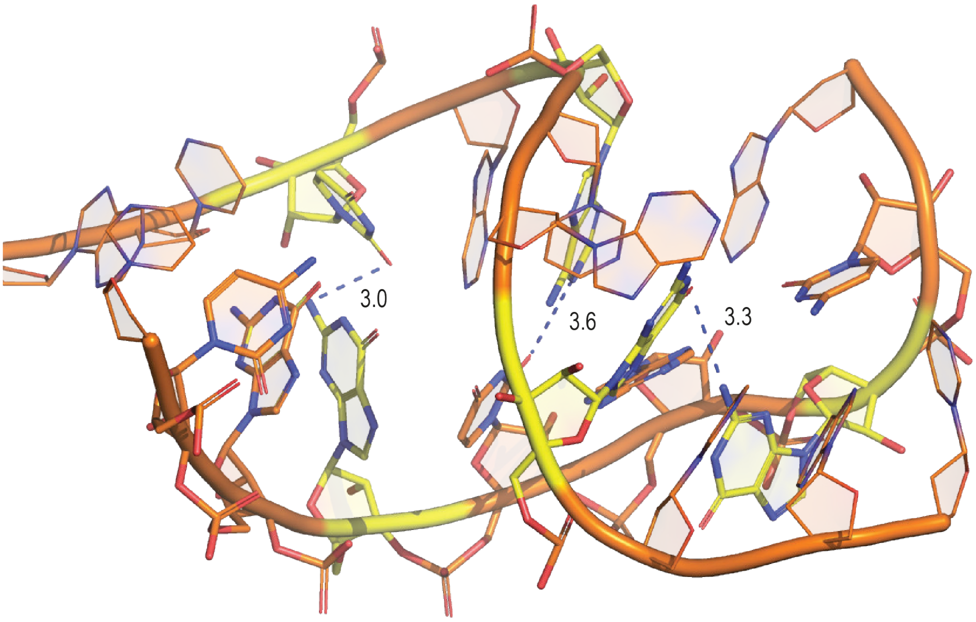

**Supplemental Figure S14. Interaction between two fragments of two RNA molecules.**

Fragments of two RNA molecules interacting around a 2-fold axis form three hydrogen bonds between G14-U'11, U15-A'9 and A16-G'6.

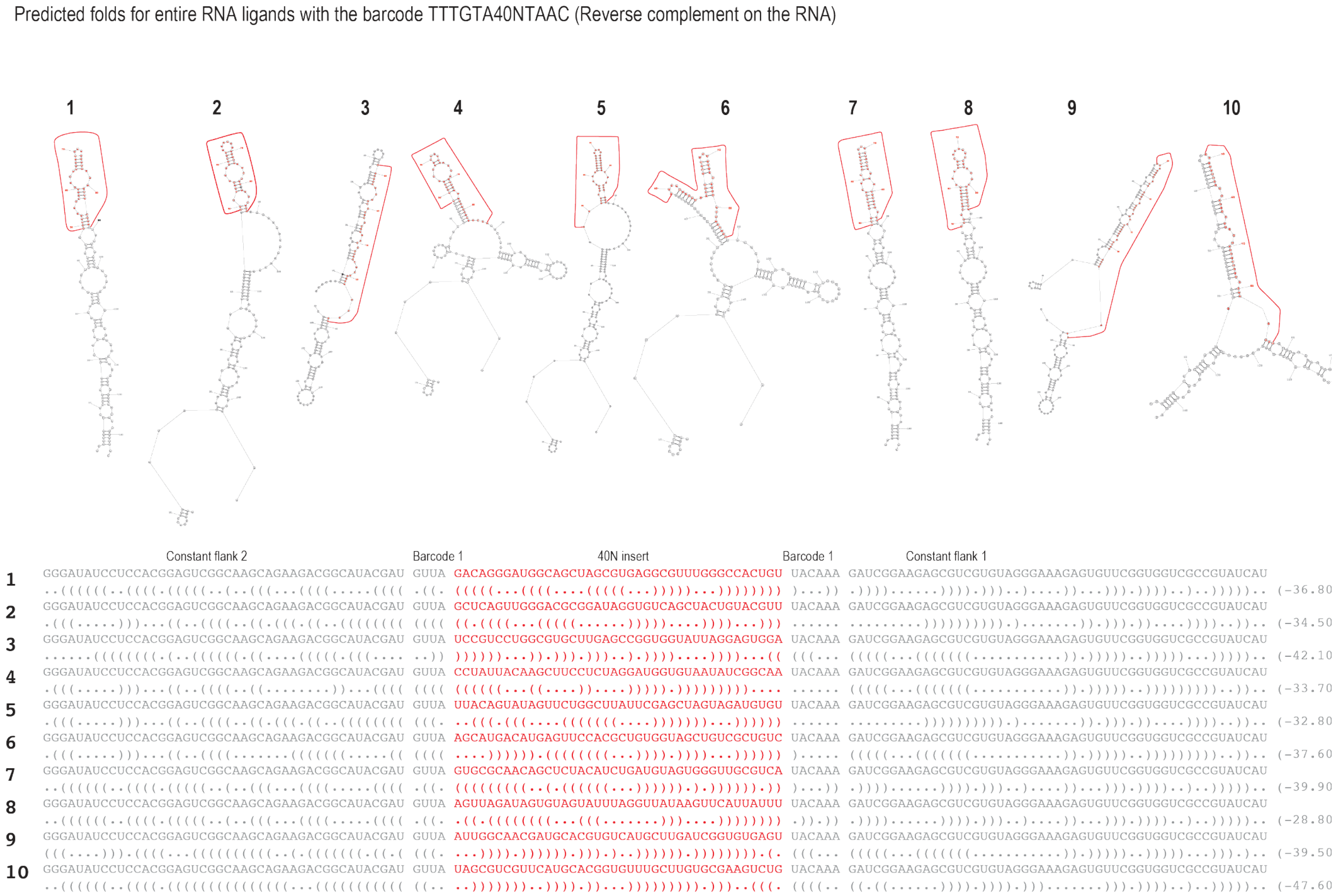

**Supplemental Figure S15. Predicted RNA secondary structures for 10 full-length selection ligands that include a random sequence flanked by the constant linker sequences.**

Sequences corresponding to the RNA selection ligands generated from the first ten sequences for barcode TTTGTA-40N-TAAC (SRA Accession PRJEB25907). Minimal energy secondary structures were predicted with the program RNAfold for each of the sequences, which are shown as structural diagrams (above) and as dot-bracket annotated sequences. Parentheses and dots indicate double and single stranded regions, respectively. The random 40 base region is indicated in red typeface and red lines. Note that the constant regions do not impose a strict bias towards particular structure for the random region.

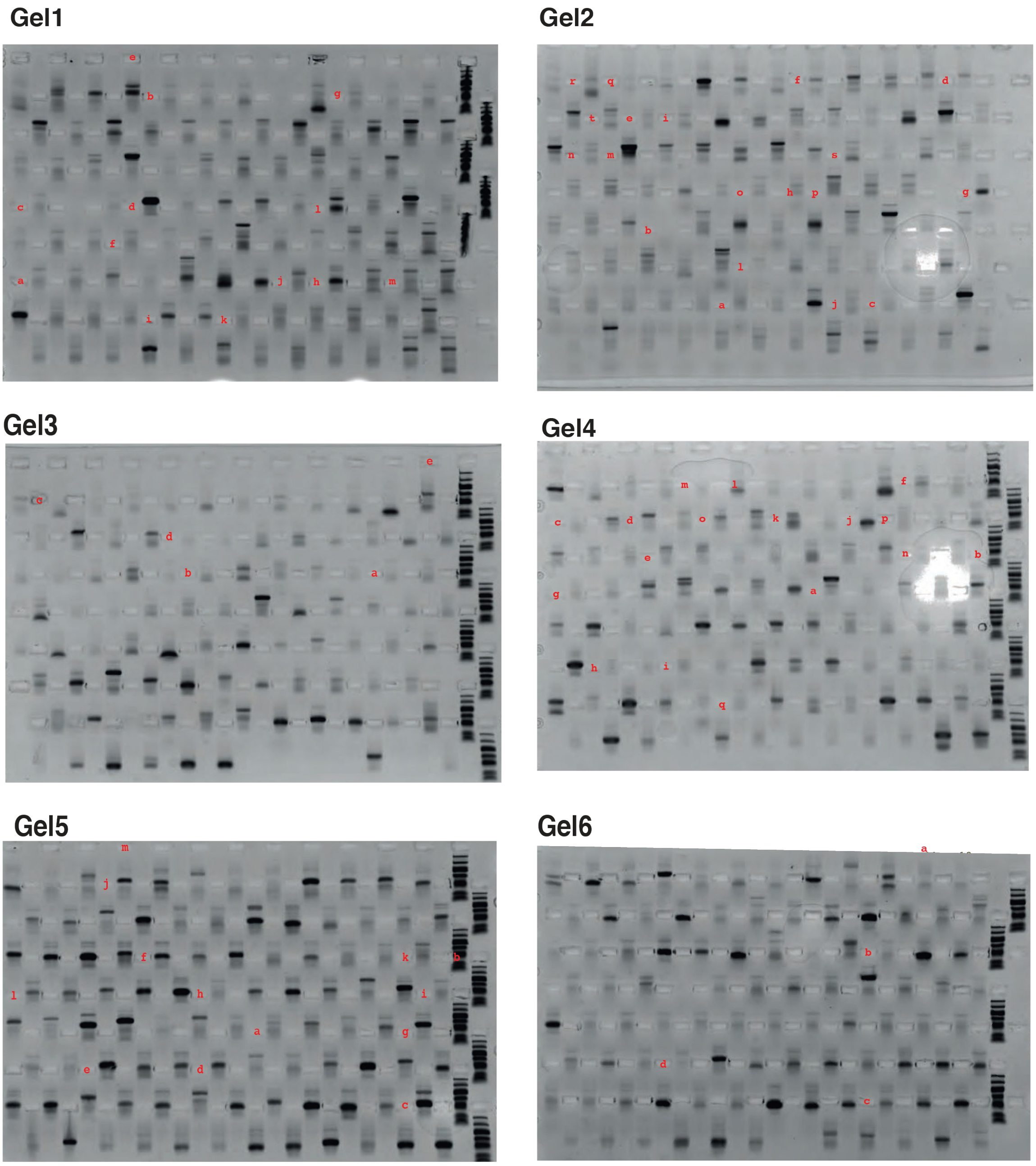

**Supplemental Figure S16. SDS page analysis of the proteins subjected to HTR-SELEX.** RBP fusion proteins were expressed in 96-well plates, purified and analyzed using 96-well SDS-PAGE gels (ePAGE, Invitrogen, run downward). Lanes containing proteins that correspond to generated motifs (see **Supplemental Table S1**) are indicated in red letters in the respective loading wells.

**Supplemental Figure 17. Annotation of RBDs in the constructs and full length protein sequences.** SMART database was used to annotate the RBDs in both constructs and full length amino acid sequences of the longest protein-coding transcripts obtained from ENSEMBL (version 99). For each construct, the full length protein is shown (top) with the aligned construct sequence (bottom). The RBDs are indicated by the colored boxes and the entire amino acid sequence is presented as a grey bar. The primary motif and secondary motif are shown on the middle and right columns, respectively (see the enclosed Supplemental_Fig_S17.html.zip).

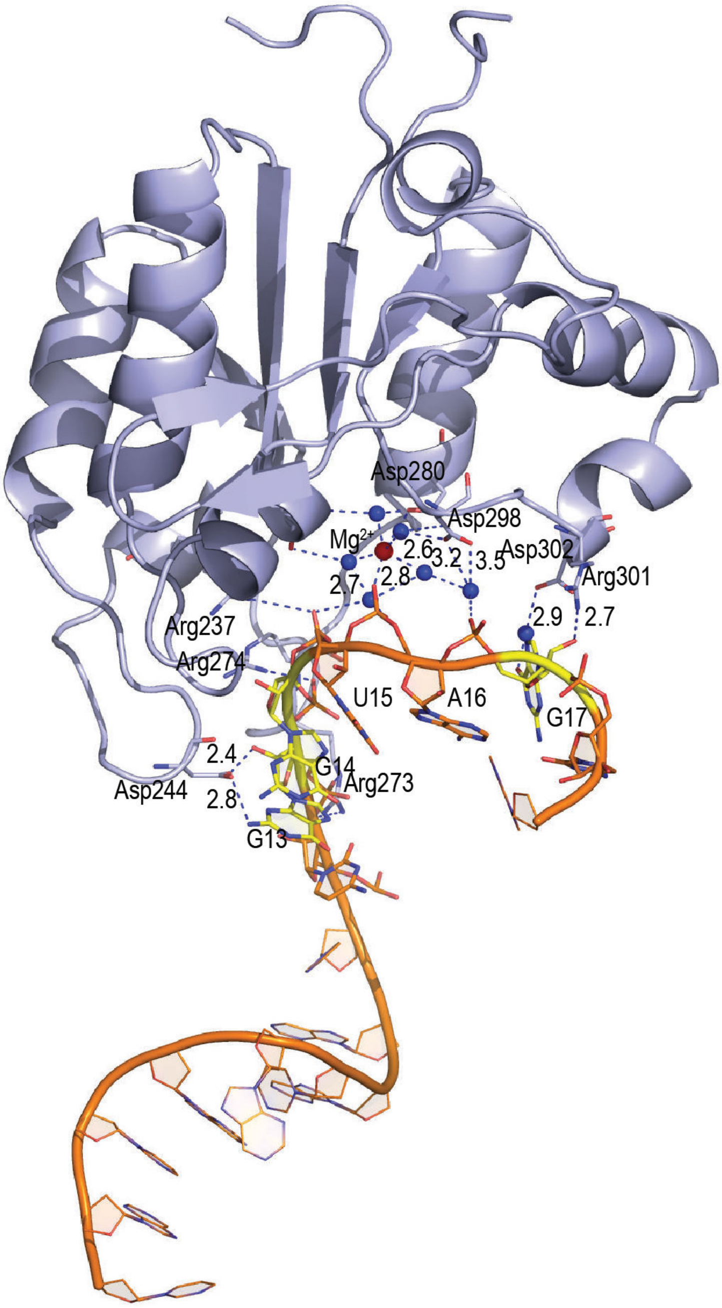

**Supplemental Figure S18. Expanded view of the structure of the ZC3H12B:RNA complex.** ZC3H12B binds to the GGUAG sequence that is located close to the 3' end of the co-crystallized RNA. Interaction between the protein and RNA molecule is mediated by a $Mg^{2+}$ ion (red sphere), water molecules (blue spheres) and multiple direct hydrogen bonds between the two macromolecules. For clarity, only the water molecules found in the active site are shown, and the involved hydrogen bonds are indicated by dashed lines (numbers indicate bond length in Å).

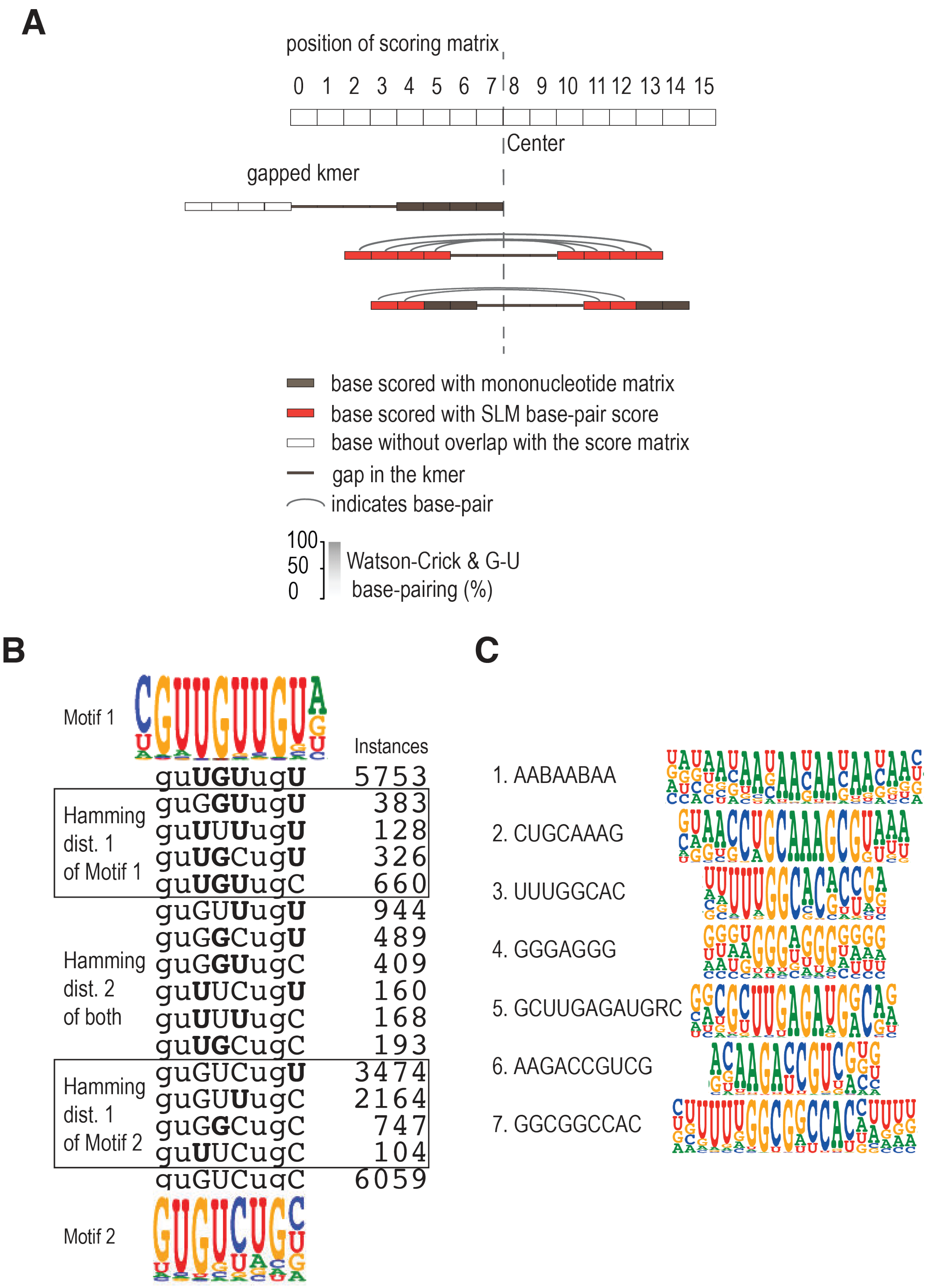

**Supplemental Figure S19. Motifs and controls**

**A)** Schematic description of the scoring process for the SLM. All possible alignment positions between an 8-mer with a 4 base gap in the middle and the model are searched in order to find the aligned position with the best score. When the 8-mer overlaps both bases of a SLM-predicted base-pair, the score for the paired position (red tiles connected by black lines) is derived from the SLM base-pair score. In cases where the kmer aligns to only one base of the SLM base-pair, the score for the position (black) is derived from the mononucleotide matrix. **B)** Seeds that represent local maxima within a Huddinge distance of one (see **Supplemental Methods**) define distinctly different motifs. Panel displays a detailed analysis of an example case of subsequence counts near seeds for LARP6 Motifs 1 and 2. The count from the fourth HTR-SELEX cycle for the consensus sequences of these two motifs, and all possible subsequences that represent the shortest edit path between them are shown. Hamming distance from the seed closer in Hamming distance to the subsequences is also indicated. Note that no subsequence in the path between the two consensus sequences has a count higher than the consensus sequences themselves. **C)** Commonly enriching background motifs. Motifs that enrich in HTR-SELEX in a large fraction of all experiments performed using unrelated *E.coli*-derived proteins are shown. These motifs represent either specific target sites for unknown RBPs derived *E. coli*, or aptamers that have affinity towards plasticware, the magnetic beads or constant parts of the fusion proteins.

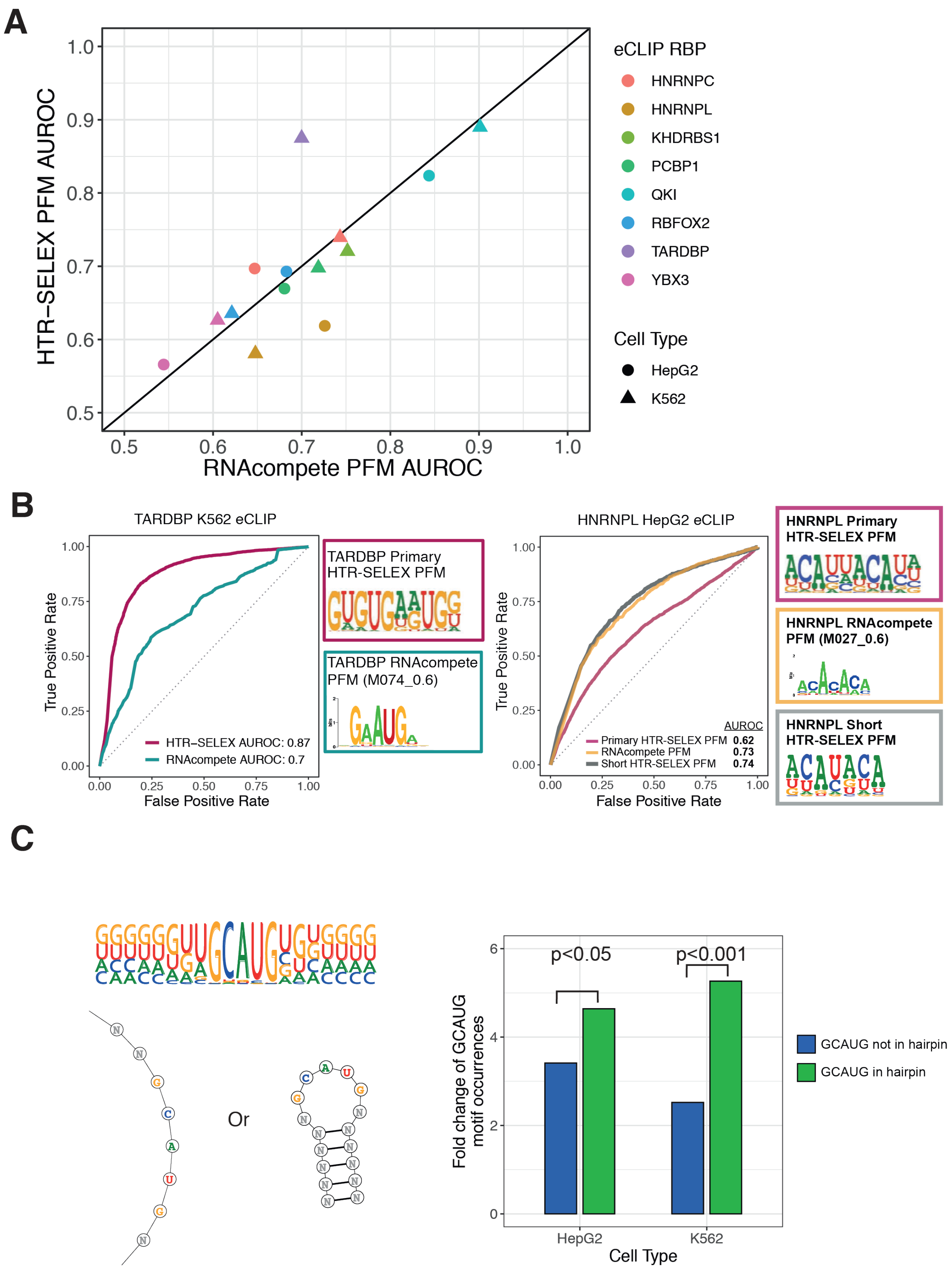

**Supplemental Figure S20. *in vivo* enrichment of the HT-SELEX motifs**

**A)** Plot compares the performance of HTR-SELEX and RNA-compete generated motifs (assessed as AUROC scores) in predicting genomic regions bound by the corresponding proteins *in vivo* based on eCLIP data. Note that the HTR-SELEX generated motif predicts *in vivo* binding better for TARDBP, whereas the RNA-compete generated motif performs better in the case of HNRNPL. **B)**

ROC plots for the two most significant outliers TARDBP and HNRNPL. HTR-SELEX motif predicts longer and higher information content motif for the TARDBP, which outperforms the short motif derived from RNAcompete. In the case of HNRNPL, our original primary HTR-SELEX motif performed worse than the RNAcompete motif. Re-analysis of the the 8-mer enrichment in the HTR-SELEX data revealed a secondary motif with similar, shorter spacing of the ACAU half-site of HNRNPL. The performance of this motif against the eCLIP data was similar to that of the RNAcompete motif. The better performance of the shorter motif over the original primary HTR-SELEX motif is potentially due to the fact that the short motif can match more than one spacing between the ACAU half-sites. **C)** Binding preference of RBFOX proteins to structured sites is confirmed by analysis of eCLIP data. Left: RBFOX1 motif and cartoons of the respective structural contexts. Right: fold change of matches to the middle GCAUG consensus in two eCLIP datasets from the indicated cell lines, compared to genomic control regions. Note that there is a larger enrichment of GCAUG matches that are within a structural context. The *p*-values for the increase in enrichment for the structured over the unstructured form are also indicated (calculated using Winflat; (Audic and Claverie 1997)).

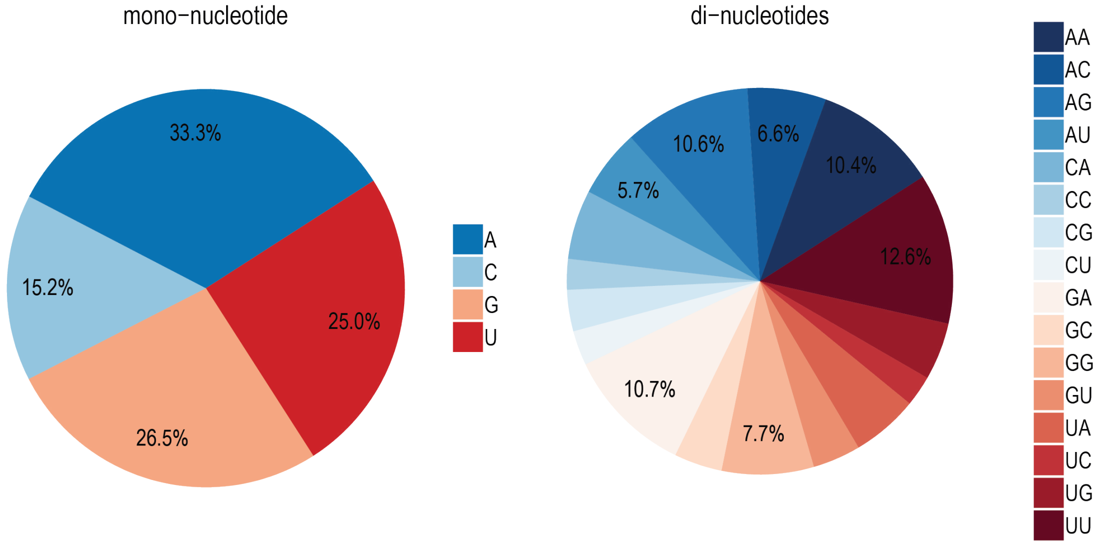

**Supplemental Figure 21. Nucleotide composition bias in the RNAcompete dataset.** Frequencies of mononucleotides (left) and dinucleotides (right) across all of the human RNAcompete motifs (downloaded from cisBP-RNA, version 0.6).

**Supplemental Table**

**Supplemental Table S1. Sequence information of proteins and DNA.**

**Supplemental Table S2. PWMs of the linear motifs.**

**Supplemental Table S3. PWMs of the structured motifs.**

**Supplemental Table S4. Dependency matrices of paired bases for the structured motifs.**

**Supplemental Table S5. X-ray data statistics and refinement parameters.**

**Supplemental Table S6. Full data for analysis of the conservation of motif matches.**

**Supplemental Table S7. Full data of the GO enrichment analysis.**

**Supplemental Table S8. Accession numbers and details of the eCLIP data used.**

**Supplemental Data**

**Supplemental Data S1. Meta-plots of the motif match enrichment near splice donor, acceptor, TSS, start and stop codon positions (y-axis scaled separately)**

**Supplemental Data S2. Meta-plots of the motif match enrichment near splice donor, acceptor, TSS, start and stop codon positions (common y-axis scale).**

**Supplemental Data S3. Histograms of the distances between motif matches and genomic features.**

**Supplemental Data S4. Count of motif matches near the genomic features.**
