## Supplementary figures and images for "Binding specificities of human RNA binding proteins towards structured and linear RNA sequences"

### TTCACG40NAGT_AAG_NNNGGUAAGGUNN_m1_c4_short.pfm.png

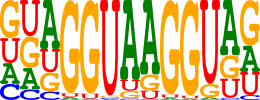

### TTCGGA40NCGC_AAG_CGGGGURUGN_m1_c3_short.pfm.png

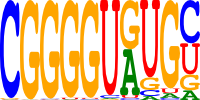

### TTCGGA40NCGC_AAG_KGUUGCGCGGG_m2_c3_short.pfm.png

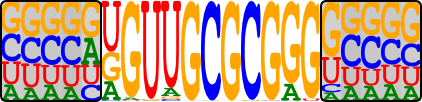

### TTCGGG40NATGT_AAG_NAGGCACR_m1_c3_short.pfm.png

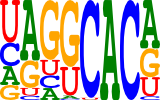

### TTCGGG40NATGT_AAG_UCACGNGCACN_m1_c3_short.pfm.png

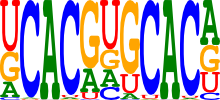

### TTCTAC40NCGA_AAG_UGUGUNUGUGU_m1_c4b-_short.pfm.png

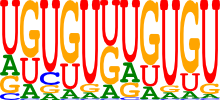

### TTCTGG40NACTT_AAG_RRGGCGUAGCGUNN_m2_c3_short.pfm.png

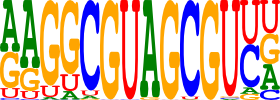

### TTCTTT40NCAG_AAG_GRGGCCGNGCGGUGG_m2_c3_short.pfm.png

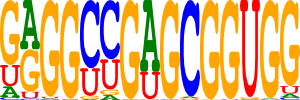

### TTCTTT40NCAG_AAG_GUGGGUCUCGG_m1_c3_short.pfm.png

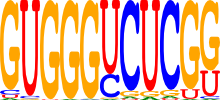

### TTGAAT40NAAGT_AAG_GCGCAUGCGNN_m1_c2_short.pfm.png

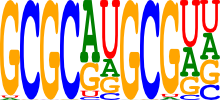

### TTGAAT40NAAGT_AAG_NGCGCAGCGUN_m1_c2_short.pfm.png

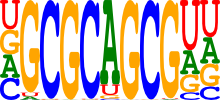

### TTGCCC40NCTC_AAG_KGUUGCGCGGG_m2_c3_short.pfm.png

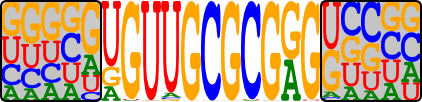

### TTGCTG40NCTA_AAG_GGGUCUUWUUUUUAA_m2_c3_short.pfm.png

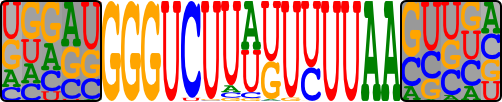

### TTGGCG40NGGGA_AAG_KUAGUWKUAGK_m1_c4_short.pfm.png

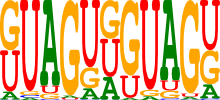

### TTGGGT40NGGC_AAH_GKDKCGACGGK_m1_c3_short.pfm.png

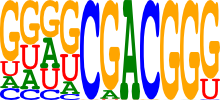

### TTGGTG40NCTAA_AAG_NGUUAGCRK_m1_c3_short.pfm.png

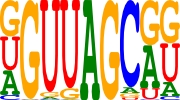

### TTGTCT40NGCT_AAG_GUWGCAUGUG_m1_c3_short.pfm.png

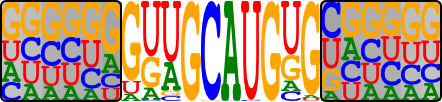

### TTTACT40NTACC_AAG_GUAGUGDGUAGK_m1_c4b-_short.pfm.png

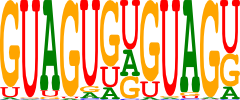

### TTTACT40NTACC_AAG_GUAGUNNNGUAGK_m1_c4b-_short.pfm.png

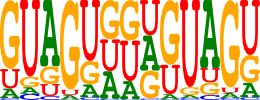

### TTTGAC40NGGT_EMO_NUGCAUAAUGN_m1_c4b0_short.pfm.png

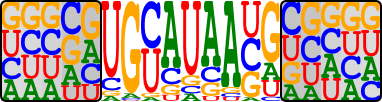

### TTTGAC40NGGT_EMO_SAUGYAURAKKUG_m2_c4b0_short.pfm.png

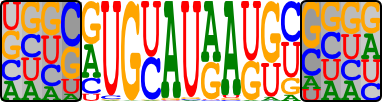

### TTTGTA40NTAAC_AAG_GUCUUGGCNNNNNNGAGCYNN_m2_c4b-_short.pfm.png

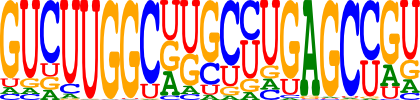

### TTTGTA40NTAAC_AAG_GUGUCNNNNNNNNNGAAGUCN_m2_c4b-_short.pfm.png

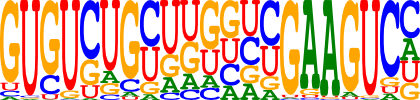

### TTTGTT40NTTAG_AAG_NGCAGUUUUAGGGCN_m2_c4_short.pfm.png

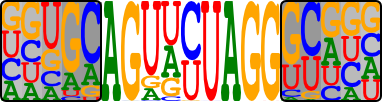
