## Supplementary figures and images for "Binding specificities of human RNA binding proteins towards structured and linear RNA sequences"

### TAAACAG26N_EMO_GCUUGC_m1_c4b0_short.pfm.png

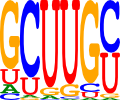

### TAAGGC40NTGCC_AAG_GUUUUUG_m1_c4_short.pfm.png

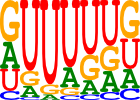

### TAAGGG40NACC_EMJ_NUGUAAAUAG_m1_c4b0_short.pfm.png

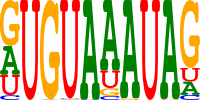

### TAAGTT40NTCGA_AAG_NUACUAACAWN_m1_c3b-_short.pfm.png

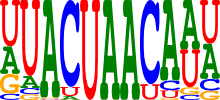

### TACATC40NCAAG_AAH_CGCCAGUGCG_m1_c3b-_short.pfm.png

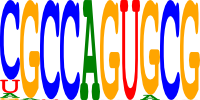

### TACCGG40NAGT_AAG_UUUUUGGUGG_m1_c4_short.pfm.png

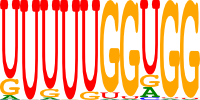

### TACCTC40NATT_AAG_GCGCAAGAGC_m1_c4_short.pfm.png

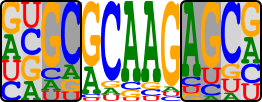

### TACCTC40NATT_AAG_NGCCCUCAAGGYN_m1_c4_short.pfm.png

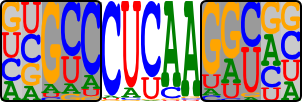

### TACCTG40NACG_AAG_NUAKAAAKUURCN_m1_c4_short.pfm.png

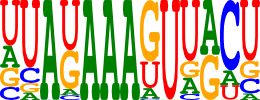

### TACCTG40NACG_AAG_NUAKAAAUUACUACAN_m2_c4_short.pfm.png

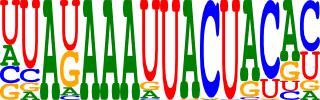

### TACGCG40NCCG_EMO_GRUCGUGAUGN_m1_c4b0_short.pfm.png

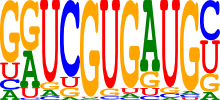

### TAGACC40NGTTA_AAG_GUAUUAAGGG_m1_c2_short.pfm.png

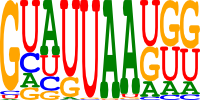

### TAGACC40NGTTA_AAG_GUGGACUUAANNN_m1_c2_short.pfm.png

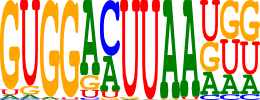

### TAGATG40NTAT_EMJ_NGGAACUAAN_m1_c4b0_short.pfm.png

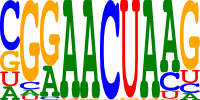

### TAGCAT40NGCAA_AAG_UAAUAGN_m1_c4_short.pfm.png

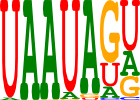

### TAGCCA40NTAT_AAG_GUUUUUUGGG_m1_c3_short.pfm.png

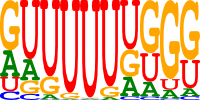

### TATAAA40NACG_AAG_GCAURCAYCC_m1_c3_short.pfm.png

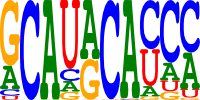

### TATGAC40NTCA_AAG_NCAGGUAAGUGN_m2_c3_short.pfm.png

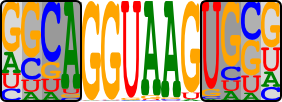

### TATGAC40NTCA_AAG_NUGCGNCAGUCGGUAGC_m2_c4_short.pfm.png

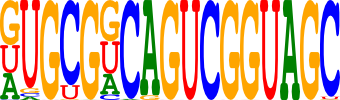

### TATGAG40NCGT_AAG_ACAUWWCAUW_m1_c3_short.pfm.png

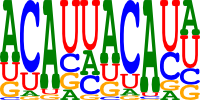

### TATGAG40NCGTAAG3_HNRNPL_ACAYACA.pfm.png

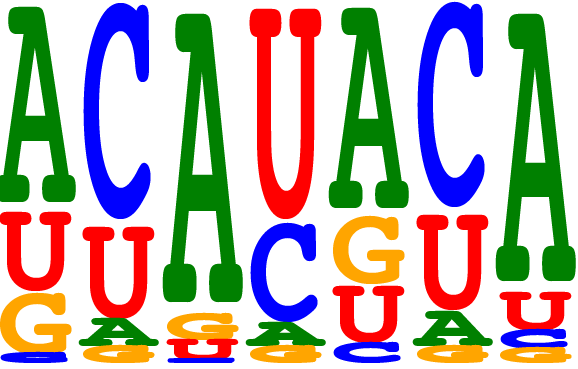

### TATGGA40NAGCC_AAG_GGKUGCGUCG_m1_c4_short.pfm.png

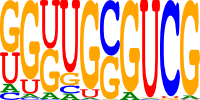

### TATGGT40NATA_EMQ_CGNUAUAUAG_m1_c4b-_short.pfm.png

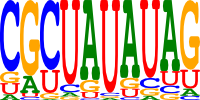

### TATGGT40NATA_EMQ_NUGCAUAAUGN_m2_c4b-_short.pfm.png

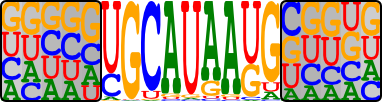

### TATTCA40NTCAG_AAG_SGUGCAUAAKGCN_m2_c3_short.pfm.png

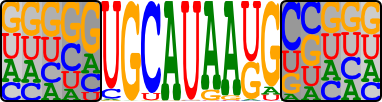

### TATTCA40NTCAG_AAG_UUAUAAUCAAC_m2_c3_short.pfm.png

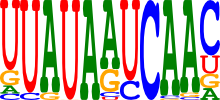

### TATTCT40NCTC_AAG_UCAUUCAU_m1_c4_short.pfm.png

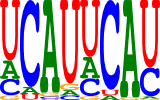

### TCAAAG40NGAG_AAG_KUAUUUAUGGUGG_m2_c3_short.pfm.png

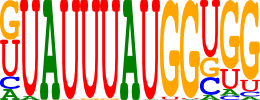

### TCAAAG40NGAG_AAG_UUAUUUAUU_m1_c2_short.pfm.png

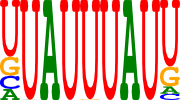

### TCACGG40NTGT_EMO_UUUUAUUUUG_m1_c4b0_short.pfm.png

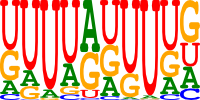

### TCAGTA40NACC_AAG_GUGUGAAUGG_m1_c4_short.pfm.png

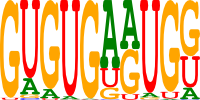

### TCAGTA40NACC_AAG_GUGUGNGUGUG_m1_c4_short.pfm.png

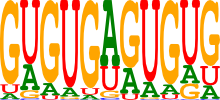

### TCATAG40NGGT_AAG_NGCCCUCAAGGYN_m1_c4b-_short.pfm.png

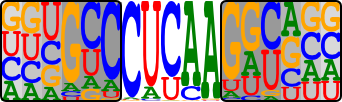

### TCATAG40NGGT_AAG_NGCGUCGUGGCN_m1_c4b-_short.pfm.png

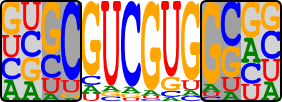

### TCCCAA40NGCGC_AAG_KUGUUGUGUUKK_m1_c3_short.pfm.png

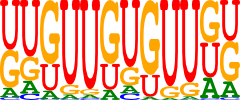

### TCCGCG40NTCT_AAG_NNUGUAYAGK_m1_c2_short.pfm.png

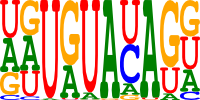

### TCCGCG40NTCT_AAG_NUGUAAAUAGKN_m2_c2_short.pfm.png

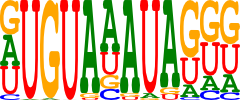

### TCCTCG40NCAG_AAG_UAAANNNNNNNUAAA_m1_c4_short.pfm.png

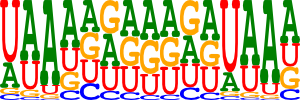

### TCCTGC40NTGCC_AAG_UUCGUGGCUUG_m1_c3_short.pfm.png

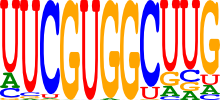

### TCCTGC40NTGCC_AAG_WGCGCGGUYUU_m1_c3_short.pfm.png

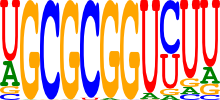

### TCCTTG40NTGGC_AAG_GGCACGGAUG_m1_c4_short.pfm.png

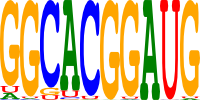

### TCCTTG40NTGGC_AAG_UUGCCGGAUGYCGU_m2_c4_short.pfm.png

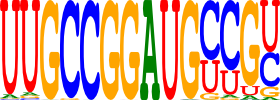

### TCGATT40NTTG_AAG_NYGUAAAKUGUAGN_m2_c4_short.pfm.png

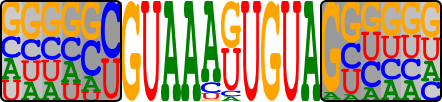

### TCGTAA40NCGG_AAG_UCCUUNNCCUU_m1_c3_short.pfm.png

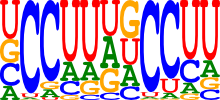

### TCGTAG40NCAGC_AAG_UAGGUUAGGG_m1_c4_short.pfm.png

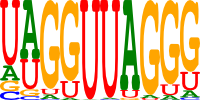

### TCGTCC40NGCC_AAG_KKGUGUGUGUGUKK_m1_c4_short.pfm.png

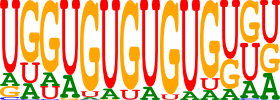

### TCGTCT40NGGTG_AAG_UNURUUUAUU_m1_c4_short.pfm.png

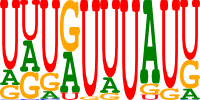

### TCTAAA40NCGC_EMN_UCACRUGCACA_m1_c4_short.pfm.png

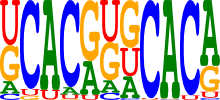

### TCTAAA40NCGC_EMN_UGCACA_m1_c4_short.pfm.png

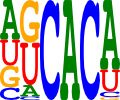

### TCTAGA40NCGA_AAG_GUUGUUGUUG_m1_c4_short.pfm.png

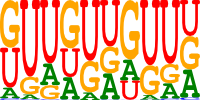

### TCTAGA40NCGA_AAG_NNUGYGUGCGUUUCCGSN_m2_c4_short.pfm.png

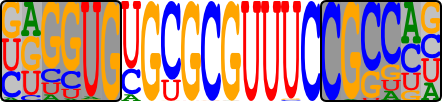

### TCTCAC40NACCT_AAG_GCCAUNNNAUCGC_m1_c4b-_short.pfm.png

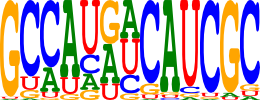

### TCTCAC40NACCT_AAG_NUAACAUCNN_m1_c4_short.pfm.png

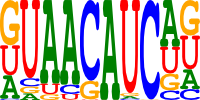

### TCTCGA40NCAT_AAG_CGUAGCGUCU_m1_c3b0_short.pfm.png

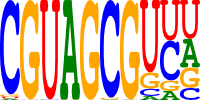

### TCTCGT40NGCC_AAH_GNUCANGNNG_m1_c3_short.pfm.png

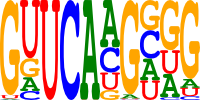

### TCTCGT40NGCC_AAH_GUUCAAGGUG_m1_c3_short.pfm.png

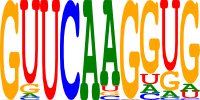

### TCTGGT40NGGG_AAH_GYUCUAGUUAGYAUC_m2_c2_short.pfm.png

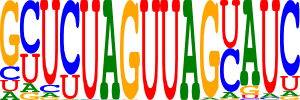

### TCTGGT40NGGG_AAH_NNUGUGUGSGUUUCCGSN_m2_c2_short.pfm.png

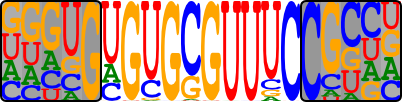

### TCTTAA40NGCA_AAG_CAGNUWCA_m1_c4_short.pfm.png

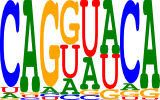

### TCTTAA40NGCA_AAH_NGCGACGGKN_m1_c3_short.pfm.png

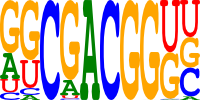

### TCTTGC40NATAG_AAG_NUAACAUCGGG_m1_c4_short.pfm.png

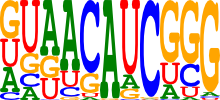

### TCTTGC40NATAG_AAG_RCCAYRHCAUCGY_m1_c4_short.pfm.png

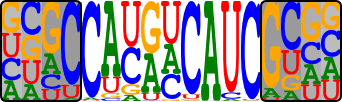

### TCTTGC40NTTC_AAG_CCUCUNUGAGG_m1_c2_short.pfm.png

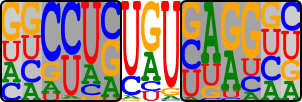

### TCTTGC40NTTC_AAG_CURCGUUUUAGKAG_m2_c2_short.pfm.png

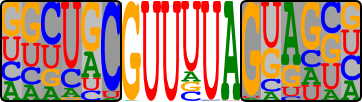

### TGAAGA40NACAG_AAH_NNUGUGUGSGUUUCCGSN_m2_c3_short.pfm.png

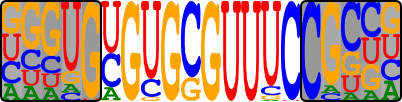

### TGAAGA40NATG_AAG_GUGUGNGUGUG_m1_c4_short.pfm.png

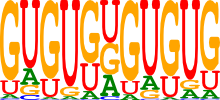

### TGAAGC40NCTA_AAG_UAACAUCA_m1_c4_short.pfm.png

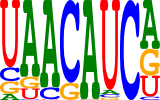

### TGAATC40NGTA_AAG_NUAGGUUAGGGW_m2_c4_short.pfm.png

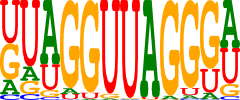

### TGAATG40NGTAG_AAG_NGCAGGUAAGUGSN_m2_c2_short.pfm.png

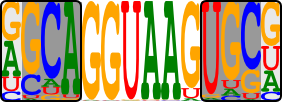

### TGAATG40NGTAG_AAG_NGCGNCAGUCGGUAGCN_m2_c3_short.pfm.png

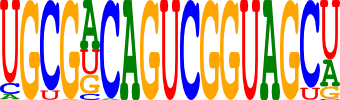

### TGATGA40NCATT_AAG_NGCAGGUAAGUGCN_m2_c4_short.pfm.png

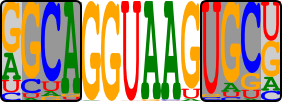

### TGATGA40NCATT_AAG_NUUGCCAWAGUKU_m2_c4_short.pfm.png

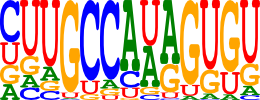

### TGCAAG40NAAC_AAG_UUAGGUUAGG_m1_c4_short.pfm.png

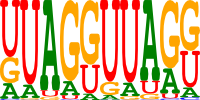

### TGCAGT40NAAG_AAG_GCCAUGAUCAUGAU_m2_c3_short.pfm.png

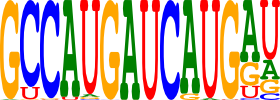

### TGCAGT40NAAG_AAG_UUGCCUUGUCCCGAU_m2_c3_short.pfm.png

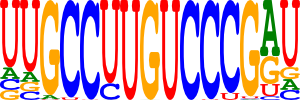

### TGCGCT40NTGA_AAG_GCAGUKUUAGGGY_m1_c3_short.pfm.png

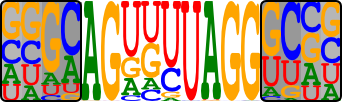

### TGCGCT40NTGA_AAG_GGGUSUACGGGGU_m2_c3_short.pfm.png

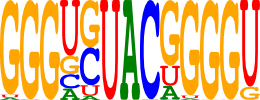

### TGCTAA40NAACA_AAG_YRCGUUUUAGKR_m1_c4_short.pfm.png

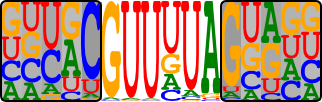

### TGGACT40NAAT_EMJ_UUUAUGNUUU_m1_c4b0_short.pfm.png

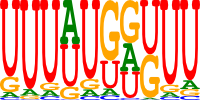

### TGGCCT40NATTT_AAG_NNUGUGUGSGUUUCCGSN_m2_c3_short.pfm.png

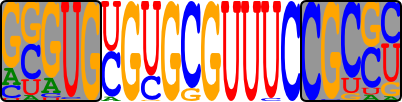

### TGGGAG40NAGG_EMO_GUGAUUG_m1_c4b0_short.pfm.png

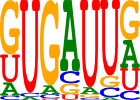

### TGGGCT40NGTT_AAG_GGUGUGGUGU_m1_c4_short.pfm.png

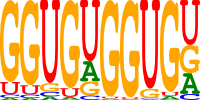

### TGGTCA40NGTG_AAG_GGGUGUGCAC_m1_c3_short.pfm.png

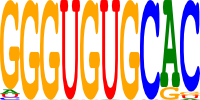

### TGGTCA40NGTG_AAG_NNCAUUGCACN_m1_c3_short.pfm.png

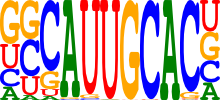

### TGGTCG40NTCGG_AAG_GKUGCAUGUG_m1_c2_short.pfm.png

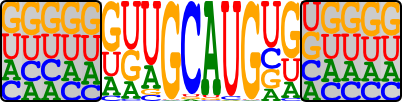

### TGGTGG40NTAA_EMO_NAGCUUNGUGCUN_m1_c4b0_short.pfm.png

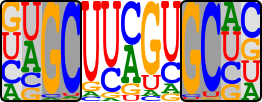

### TGGTGG40NTAA_EMO_NGCUUCGNGCN_m1_c4b0_short.pfm.png

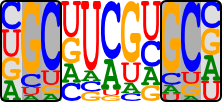

### TGTATG40NCGG_AAG_NNAGAGUUUAUNN_m1_c3_short.pfm.png

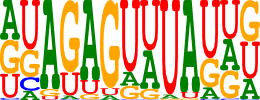

### TGTCGA40NCCGA_AAH_KUAGKUUAGGK_m1_c3_short.pfm.png

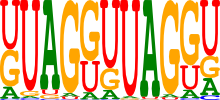

### TGTCGA40NCCGA_AAH_KUAGKUUUAGGK_m1_c3_short.pfm.png

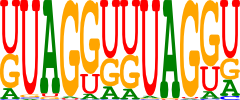

### TGTCTA40NTCG_AAG_CAUUGCACN_m1_c2_short.pfm.png

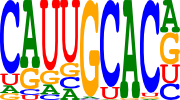

### TGTCTA40NTCG_AAG_NUGCACR_m1_c2_short.pfm.png

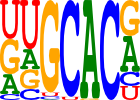

### TGTCTC40NTCA_AAG_UAGAGUUUAU_m1_c4_short.pfm.png

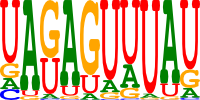

### TGTCTC40NTCA_AAG_UAUUUAU_m1_c4_short.pfm.png

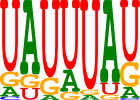

### TGTGGT40NGGC_EMJ_UAAACAAUAA_m1_c4b0_short.pfm.png

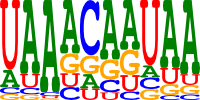

### TGTTCG40NGAC_AAG_UGUGUNUGUGU_m1_c4_short.pfm.png

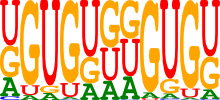

### TTATGA40NCAG_AAG_YRCGCUCCUNGCY_m2_c2_short.pfm.png

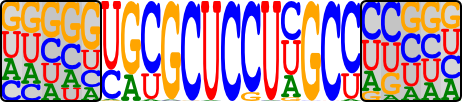

### TTATGA40NCAG_AAG_YRYGCUCGUGCYN_m2_c2_short.pfm.png

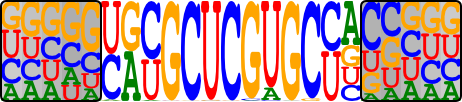

### TTATGG40NCGTT_AAH_UGUGUUGUGU_m1_c3_short.pfm.png

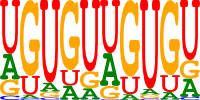

### TTCACG40NAGT_AAG_NNGGUAGGURRN_m1_c4_short.pfm.png

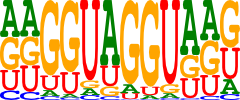
